# GeneFamHunter: annotation-independent recovery of divergent and tandemly arrayed gene-family members from genome assemblies

**DOI:** 10.64898/2026.11.02.763643

**Authors:** Pavel B. Klimov, Qixin He

## Abstract

**Motivation:** Gene families expand by tandem duplication, diverge and decay, so their members are often missing, fragmented or fused in automated genome annotations, and divergent copies escape pairwise similarity searches. These challenges are particularly pronounced in non-model organisms, for which well-annotated genomes and reference proteomes are often unavailable, limiting the applicability of protein-based gene family recovery methods.

**Results:** We developed GeneFamHunter to recover gene-family members directly from nucleotide assemblies. It searches continuous six-frame translations of overlapping genomic windows using four complementary domain- and profile-based methods, clusters hits while distinguishing tandem paralogs, predicts and validates gene models, and constructs gene trees. Across six Quest for Orthologs genomes, window-based HMMER searches recovered all 130 reference papain-like (C1A) loci, with 21 off-target hits, compared with 45 for stop-to-stop ORF searches; whole-contig HMMER searches recovered 108 (83.1%). With 11 distantly related queries, PSI-BLAST and DELTA-BLAST recovered all reference loci, whereas RPSTBLASTN recovered 98.5–99.2% and TBLASTN, as used by BITACORA, recovered 94.6– 95.4%. Across six mite genomes, GeneFamHunter recovered all 209 previously reported C1 loci and identified 37 additional candidate loci. In a separate comparison, it recovered all 128 reference UniProt C1A proteins and identified five additional loci without annotated gene models; OrthoFinder included 35 non-C1A proteins, whereas OMA recovered 98 reference proteins.

**Availability and implementation:** GeneFamHunter is implemented in Bash for Linux and SLURM clusters and is freely available at https://github.itap.purdue.edu/HeLab/GeneFamHunter (MIT license). Benchmark code and data: https://github.itap.purdue.edu/HeLab/GeneFamHunter_benchmarks and Zenodo (DOI: 10.5281/zenodo.23744704).

**Supplementary information:** Supplementary data are available at *Bioinformatics* online.

## 1 Introduction

Expansion, contraction and divergence of gene families drive much of genome evolution, so delineating family members is a routine first step in comparative genomics (Nei and Rooney, 2005; Demuth and Hahn, 2009). Directly inferring gene families from genome assemblies is difficult for three reasons.

Members of rapidly evolving families may share only ~25% amino-acid identity, in the ‘twilight zone’ where pairwise similarity searches lose sensitivity (Rost, 1999). Recent paralogs often lie in dense tandem arrays that gene predictors merge into chimeric models. Annotations of non-model genomes are frequently incomplete, fragmented or missing (Denton *et al*., 2014; Salzberg, 2019), and annotation quality alone can change orthology-inference performance by up to 30% (Trachana *et al*., 2011).

Existing tools address parts of this problem. Orthology inference methods such as OrthoFinder (Emms and Kelly, 2019), OMA (Altenhoff *et al*., 2019), InParanoid (Östlund *et al*., 2010) and OrthoMCL (Li *et al*., 2003) start from predicted proteomes and inherit their errors. Pipelines using unannotated genomes, such as BITACORA (Vizueta *et al*., 2020) avoid this dependence but rely on TBLASTN, which loses sensitivity when only distant homologs are available as queries. TGFam-Finder (Kim *et al*., 2020) searches profile HMMs against six-frame genomic translations, but it splits translations at stop codons, which in eukaryotes often lie in introns, and it does not explicitly separate closely spaced tandem paralogs.

Here, we introduce GeneFamHunter, which infers gene families directly from raw genome assemblies. It searches continuous six-frame translations of overlapping genomic windows with four complementary methods, clusters hits with explicit rules that keep tandem paralogs apart, excludes fused proteins from the evidence, and validates predicted proteins before gene-tree inference (Fig. 1A). Across three benchmarks, GeneFamHunter achieved 100% recall of the reference gene-family members while identifying additional candidate loci, demonstrating its ability to recover family members directly from genome assemblies without relying on existing gene annotations.

**Fig 1.**
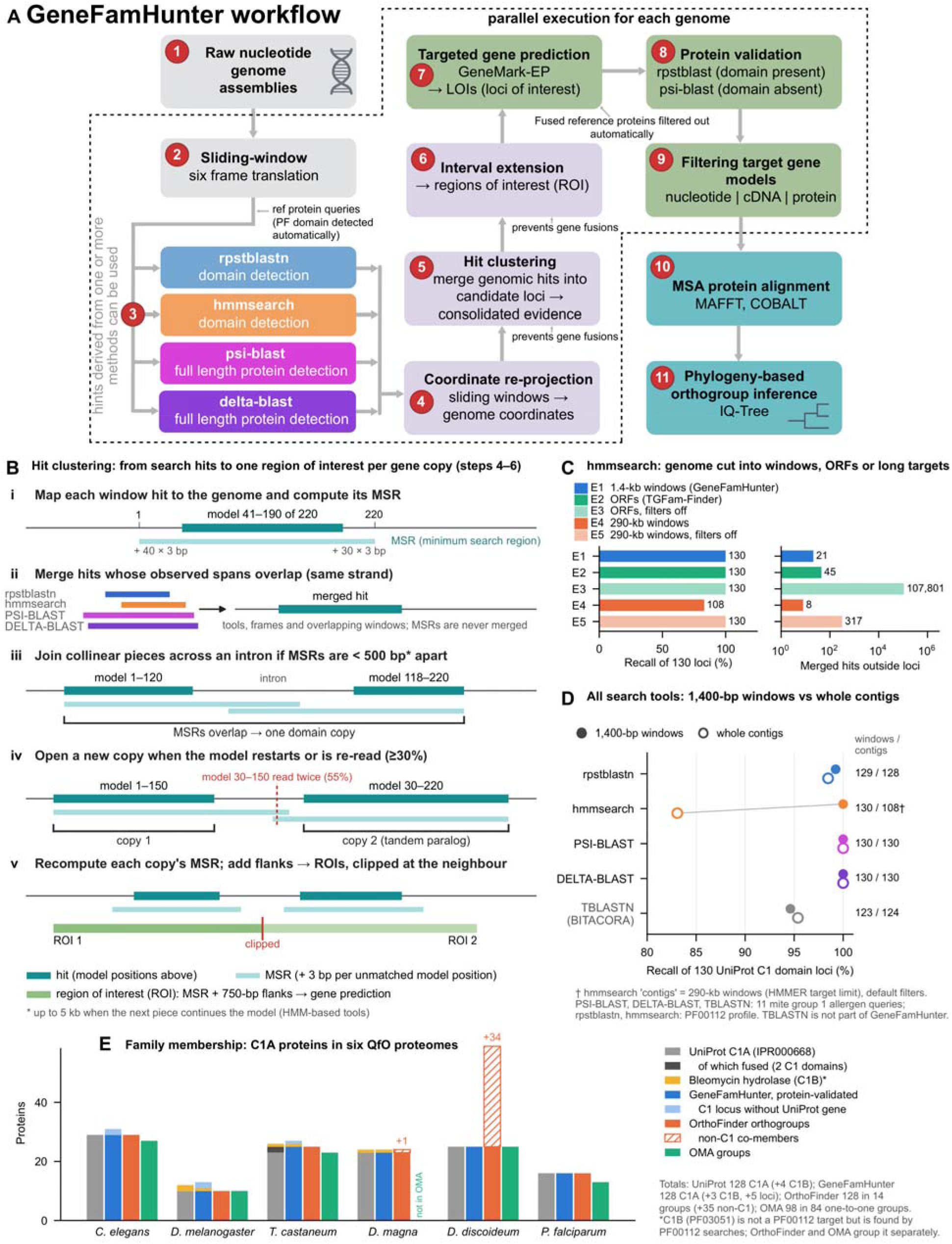
GeneFamHunter workflow and benchmark. (**A**) Workflow; steps 2–9 run in parallel for each genome. (**B**) Hit clustering into one region of interest per gene copy (steps 4–6). Hits (teal; model positions above) are shown to scale with their minimum search regions (MSRs): (i) each hit is extended to its MSR; (ii) overlapping observed hits are merged, MSRs never; (iii) pieces of one domain are joined across an intron when their MSRs are < 500 bp apart; (iv) a hit that restarts or re-reads ≥ 30% of the model opens a new copy, even when MSRs overlap; (v) each copy’s MSR plus flanks forms a region of interest, clipped at the neighbour. (**C, D**) Searches of six Quest for Orthologs genomes scored against 130 UniProt C1A domain loci (128 proteins). (**C**) hmmsearch recall and merged hits outside reference loci. E1, GeneFamHunter’s 1,400-bp windows; E2, stop-to-stop ORFs as in TGFam-Finder; E4, 290-kb windows; E3/E5 (lighter), acceleration filters off. (**D**) The four GeneFamHunter search tools and TBLASTN (the BITACORA search; not part of GeneFamHunter) on 1,400-bp windows (filled) and whole contigs (open; for hmmsearch, 290-kb windows, E4); numbers are loci recovered. PSI-BLAST, DELTA-BLAST and TBLASTN used the same 11 mite group 1 allergen queries. (**E**) C1A family membership in the six QfO proteomes (Supplementary Note S6): UniProt C1A proteins (InterPro IPR000668; dark, fused models with two C1 domains), GeneFamHunter after protein-level validation (light, C1 loci without an annotated gene), and C1A proteins in OrthoFinder orthogroups and OMA groups (hatched, co-members without a C1 domain). Yellow, bleomycin hydrolases (C1B, PF03051): not a PF00112 target but found by PF00112 searches; OrthoFinder and OMA place them in separate groups. *D. magna* is not in OMA.

## 2 Implementation

### 2.1 Input and search

GeneFamHunter is implemented as a set of Bash scripts that support two execution modes: a standalone mode that processes up to two genomes concurrently and a SLURM-based mode that processes genomes in parallel using job arrays. Both modes are followed by a joint gene-tree construction step. Inputs are nucleotide assemblies of any contiguity, at least one protein query from the target family, and optional protein evidence. The target domain is identified automatically as the best RPS-BLAST hit of the queries in the Conserved Domain Database (CDD; Wang *et al*., 2023).

Each assembly is cut into overlapping windows (default 1,400 bp with a 700-bp step; BEDTools, Quinlan and Hall, 2010) that are translated in all six frames without breaking at stop codons (SeqKit; Shen *et al*., 2016). Continuous translation lets a domain be detected across an intron that contains an in-frame stop codon, and short targets keep every sequence below HMMER’s target-length limit (~100,000 residues).

The windows are searched by four methods (Fig. 1A, step 3): RPSTBLASTN (BLAST+; Camacho *et al*., 2009) against the target-domain profile, with the database size set to that of the whole CDD (E ≤ 10□□); hmmsearch with the Pfam HMM (HMMER 3.3.2, Eddy, 2011; Pfam, Mistry *et al*., 2021; domain i-E-value < 0.01); PSI-BLAST with the protein queries (Altschul *et al*., 1997; three iterations, E ≤ 10□^3^); and DELTA-BLAST, which seeds its profile with conserved domains (Boratyn *et al*., 2012).

### 2.2 Clustering hits and resolving tandem paralogs

Hits are clustered per tool, reference model, contig and strand in transcription order, using rules designed for gene families in which each gene contains a single copy of the selected target domain (ie., gene family-defining domain), although it may contain other, distinct domains (Fig. 1B). (i) Each hit is first extended to its minimum search region (MSR), the genomic span the full domain would occupy without introns (3 bp per unmatched model position). (ii) Hits whose observed genomic spans overlap, including duplicates from overlapping windows and frames, are merged; MSRs are never merged, because those of neighbouring tandem copies overlap by design. (iii) A hit that continues the model where the previous piece ended (model positions advancing) is joined to it as part of the same domain interrupted by an intron, provided that the gap between their MSRs is < 500 bp, or < 5 kb for HMM-based hits when the new piece starts within −9 to +30 model positions of where the previous one stopped. (iv) A hit that restarts the model, does not extend it, or re-reads ≥ 30% of it opens a new domain copy; this separates tandem paralogs even when they are only a few hundred base pairs apart. The MSR of each copy is then recomputed from all of its pieces. Copies from different tools are merged when their observed hits overlap on the same strand.

Each MSR is extended by flanks (750 bp for domain hits; 360/180 bp for full-length protein hits) into a region of interest (ROI). Flanks are clipped where they reach the MSR or hits of a neighbouring copy, so that each ROI is expected to contain a single gene (Fig. 1B, v).

### 2.3 Gene prediction and validation

Before gene prediction, protein evidence is screened with hmmscan, and proteins with two or more target-domain regions separated by > 20 residues are removed as probable fusions of tandem genes (Supplementary Note S1). Gene models are predicted with GeneMark-EP+ and ProtHint (Brůna *et al*., 2020), trained on the whole genome, and kept only within ROIs and only if they overlap the search hits.

Predicted proteins are searched against the CDD with RPS-BLAST (E ≤ 10□^3^); a model is confirmed if it carries a domain from the target CDD superfamily (for C1A, including C1B bleomycin hydrolases). If no domain profile exists, validation uses PSI-BLAST with the query proteins instead. The high-quality set contains confirmed models (and split fragments with the target domain) that cover ≥ 50% of the merged-hit span of their ROI. These proteins are aligned with MAFFT (Katoh and Standley, 2013) or COBALT (Papadopoulos and Agarwala, 2007), and a gene tree with SH-aLRT and ultrafast bootstrap support is inferred with IQ-TREE 2 (Minh *et al*., 2020).

### 2.4. Use of AI-assisted tools

ChatGPT (OpenAI) and Claude (Anthropic, with Opus 5.5) were used to assist with editing manuscript text and reviewing and editing source code. All AI-assisted changes were reviewed and verified by the authors, who take full responsibility for the final manuscript and software.

## 3 Benchmarks and application

We used the papain-like cysteine proteases (peptidase C1A, PF00112), which include the group 1 allergens of house dust mites and form tandem arrays in many genomes. Because gene-model accuracy depends on the external predictor, Section 3.1 evaluates the search and clustering steps, Section 3.2 family delineation, and Section 3.4 the complete pipeline.

### 3.1 Window-based searches

The reference comprised 130 C1A domain loci of 128 proteins (InterPro IPR000668), projected from UniProt annotations onto six phylogenetically diverse Quest for Orthologs genomes (Altenhoff *et al*., 2016; Supplementary Note S2). Searches were also scored against an RPSTBLASTN-based reference (Supplementary Note S2). To mimic a non-model setting, the query proteins were 11 distantly related mite group 1 allergens. We compared HMMER searches of GeneFamHunter’s 1,400-bp windows (E1) with searches of stop-to-stop ORFs, the target representation used by TGFam-Finder (E2) and of 290-kb windows approximating whole contigs (E4), with HMMER acceleration filters on (default) or off (E3, E5). The windowed search recovered all 130 loci with the fewest merged hits outside them (21, compared with 45 for ORFs; Fig. 1C). With default filters, 290-kb windows missed 22 loci (83.1% recall).

Disabling the filters restored recall but produced 317 and 107,801 off-target merged hits. Windows did not shorten hits (median model coverage 31% for both; Supplementary Fig. S3).

Unlike hmmsearch, the BLAST-based tools gave nearly identical recall on windows and whole contigs (Fig. 1D). With the same 11 distant queries, PSI-BLAST and DELTA-BLAST recovered all 130 loci, whereas TBLASTN, the search used by BITACORA, recovered 123–124 (94.6–95.4%); RPSTBLASTN with the C1 profile recovered 129 and 128 (Supplementary Table S3). At the ROI stage, all 130 reference loci were recovered without fusing adjacent genes. The same patterns held in six mite genomes, evaluated against RPSTBLASTN-supported ROIs (Supplementary Note S3).

### 3.2 Family membership compared with orthology groups

Orthology groups often serve as gene families, so we compared GeneFamHunter with OrthoFinder orthogroups and OMA groups (Altenhoff *et al*., 2024) in the six QfO proteomes (Supplementary Note S6). The reference comprised the 128 UniProt C1A proteins (InterPro IPR000668), which carry the 130 domains of Section 3.1; bleomycin hydrolases (C1B, Pfam PF03051), which PF00112 searches also find, were counted separately. Without gene prediction in the QfO runs, UniProt proteins within ROIs stood in for gene models. GeneFamHunter recovered all 128 reference proteins, three of four bleomycin hydrolases and five C1 loci without an annotated gene; validation removed 12 nucleotide-level hits, mostly propeptide-only (I29) proteins (Fig. 1E). OrthoFinder split the 128 among 14 groups, which contained 35 proteins without a C1 domain; OMA placed 98 in 84 one-to-one groups.

### 3.3 Fused gene models in public annotations

Among 190 C1A proteins from five GenBank mite proteomes, 18 (9.5%) contained two to seven PF00112 domains, compared with 2 of 128 (1.6%; both unreviewed *T. castaneum* entries) in the six QfO proteomes (Fig. 1E) and none of 256 Swiss-Prot C1A proteins. In an RNA-seq-supported example, the drop in coverage between the two domains is consistent with adjacent tandem genes joined across a mis-annotated intron (Supplementary Fig. S1). These proteins are removed from the gene-prediction evidence.

### 3.4 Application to six mite genomes

We applied the full pipeline to the six astigmatic mite genomes analysed by Shi *et al*. (2023). GeneFamHunter produced 246 high-quality C1 gene models in 245 ROIs; 242 were confirmed by the CDD check. Shi *et al*. (2023) reported 211 genomic loci of C1 genes. Two of these have no C1 domain in a CDD search (Supplementary Table S6). All 209 remaining loci overlapped GeneFamHunter hits, and

200 (95.7%) overlapped a final gene model. The other nine (*T. putrescentiae*) had ROIs but no model that passed the filters. GeneFamHunter found 39 further loci, 37 of them confirmed C1 genes (12 full-length; 25 partial models). Where both studies predicted a gene at the same locus, 128 of 198 proteins (65%) were identical or nearly identical (Supplementary Table S6).

Annotation-dependent methods, including orthology-inference tools, can only use genes present in the annotated proteome. In the automated gene annotations of the same assemblies (provided by Shi *et al*.; not published), hmmsearch found 174 C1 loci. GeneFamHunter recovered 162 of these, and 35 of its genes did not overlap any annotated gene. Twenty-eight annotated models contained 2–8 C1 domains; Shi *et al*. (2023) separated such fused models by manual curation, whereas GeneFamHunter separated them automatically into 76 genes.

MEROPS identifiers (Rawlings *et al*., 2018) mapped onto the gene tree (Supplementary Fig. S8) placed the group 1 allergen homologs in two supported clades, peptidase 1 (mite) (C01.073) and Blo t 1-type (C01.990); GeneFamHunter’s genomic coordinates show that many of these genes form tandem arrays, for example ten C01.073 copies within 20 kb in *S. scabiei*. Although only 110 of 246 models were complete, the tree agrees with MEROPS: 111 of 114 labelled proteins in supported clades share their clade’s identifier. The 51 unlabelled proteins include a supported *T. putrescentiae* clade of 16 genes (C18), 14 found only by GeneFamHunter, which may represent an uncharacterized C1 lineage, and a one-to-one ortholog group present in all six mites (C19) that is sister to cathepsin O but has no close MEROPS match (Supplementary Note S4). Each genome (55–97 Mb) was processed in 3.6–6.9 h on 64 CPU cores.

## 4 Discussion

GeneFamHunter separates three tasks that conventional annotation pipelines often combine: locating gene-family members, delimiting individual genes, and predicting exon–intron structures. OrthoFinder infers orthogroups across the entire proteome, whereas GeneFamHunter targets a specific gene family, using domain profiles to define family membership and validate candidate loci, thereby reducing false-positive assignments. In the QfO C1A benchmark, OrthoFinder recovered the same set of C1 proteins but distributed them among 14 orthogroups and also included 35 non-C1 proteins. GeneFamHunter provides an alternative to the TBLASTN-based search used by BITACORA, which missed approximately 5% of reference loci with distant queries in our benchmark. Its six-frame translation strategy also improves on stop-to-stop ORF-based searches used by TGFam-Finder, reducing off-target hits by approximately half while explicitly distinguishing tandem paralogs. The complete workflow processes raw genome assemblies into validated gene models and gene trees in a single run, without requiring manual curation.

Its independence from existing gene annotations makes it particularly useful for newly sequenced and non-model genomes.

The current workflow assumes one target domain per gene; consequently, gene families containing naturally repeated copies of the target domain cannot be processed reliably. Default parameters are optimized for genomes with relatively short introns. For genomes with longer introns, the MSR-gap limits and ROI flanks should be increased (Supplementary Table S8). Gene-model completeness also depends on the external predictor, GeneMark-EP, which relies on protein evidence rather than transcriptome evidence for gene prediction. Future versions will incorporate predictors that use transcriptome evidence, such as BRAKER (Gabriel *et al*., 2024) and GeMoMa (Keilwagen *et al*., 2016), and support the export of corrected GFF3 annotations to facilitate the correction of fused gene models in public databases.

## References

Altenhoff, A.M. et al. (2016) Standardized benchmarking in the quest for orthologs. Nat. Methods, 13, 425–430.

Altenhoff, AM, et al. (2019). OMA standalone: orthology inference among public and custom genomes and transcriptomes. Genome Res., 29, 1152–1163.

Altenhoff, A.M. et al. (2024) OMA orthology in 2024: improved prokaryote coverage, ancestral and extant GO enrichment, a revamped synteny viewer and more in the OMA Ecosystem. Nucleic Acids Res., 52, D513–D521.

Altschul, S.F. et al. (1997) Gapped BLAST and PSI-BLAST: a new generation of protein database search programs. Nucleic Acids Res., 25, 3389–3402.

Boratyn, G.M. et al. (2012) Domain enhanced lookup time accelerated BLAST. Biol. Direct, 7, 12.

Brůna, T. et al. (2020) GeneMark-EP+: eukaryotic gene prediction with self-training in the space of genes and proteins. NAR Genom. Bioinform., 2, lqaa026.

Camacho, C. et al. (2009) BLAST+: architecture and applications. BMC Bioinformatics, 10, 421.

Demuth, J.P. and Hahn, M.W. (2009) The life and death of gene families. BioEssays, 31, 29–39.

Denton, J.F. et al. (2014) Extensive error in the number of genes inferred from draft genome assemblies. PLoS Comput. Biol., 10, e1003998.

Eddy, S.R. (2011) Accelerated profile HMM searches. PLoS Comput. Biol., 7, e1002195.

Emms, D.M. and Kelly, S. (2019) OrthoFinder: phylogenetic orthology inference for comparative genomics. Genome Biol., 20, 238.

Gabriel, L. et al. (2024) BRAKER3: fully automated genome annotation using RNA-seq and protein evidence with GeneMark-ETP, AUGUSTUS, and TSEBRA. Genome Res., 34, 769–777.

Katoh, K. and Standley, D.M. (2013) MAFFT multiple sequence alignment software version 7: improvements in performance and usability. Mol. Biol. Evol., 30, 772–780.

Keilwagen, J. et al. (2016) Using intron position conservation for homology-based gene prediction. Nucleic Acids Res., 44, e89.

Kim, S. et al. (2020) TGFam-Finder: a novel solution for target-gene family annotation in plants. New Phytol., 227, 1568–1581.

Li, L. et al. (2003) OrthoMCL: identification of ortholog groups for eukaryotic genomes. Genome Res., 13, 2178–2189.

Minh, B.Q. et al. (2020) IQ-TREE 2: new models and efficient methods for phylogenetic inference in the genomic era. Mol. Biol. Evol., 37, 1530–1534.

Mistry, J. et al. (2021) Pfam: the protein families database in 2021. Nucleic Acids Res., 49, D412–D419.

Nei, M. and Rooney, A.P. (2005) Concerted and birth-and-death evolution of multigene families. Annu. Rev. Genet., 39, 121–152.

Östlund, G. et al. (2010) InParanoid 7: new algorithms and tools for eukaryotic orthology analysis. Nucleic Acids Res., 38, D196–D203.

Papadopoulos, J.S. and Agarwala, R. (2007) COBALT: constraint-based alignment tool for multiple protein sequences. Bioinformatics, 23, 1073–1079.

Quinlan, A.R. and Hall, I.M. (2010) BEDTools: a flexible suite of utilities for comparing genomic features. Bioinformatics, 26, 841–842.

Rawlings, N.D. et al. (2018) The MEROPS database of proteolytic enzymes, their substrates and inhibitors in 2017 and a comparison with peptidases in the PANTHER database. Nucleic Acids Res., 46, D624–D632.

Rost, B. (1999) Twilight zone of protein sequence alignments. Protein Eng., 12, 85–94.

Salzberg, S.L. (2019) Next-generation genome annotation: we still struggle to get it right. Genome Biol., 20, 92.

Shen, W. et al. (2016) SeqKit: a cross-platform and ultrafast toolkit for FASTA/Q file manipulation. PLoS One, 11, e0163962.

Shi, L. et al. (2023) Comparative analysis of cysteine proteases reveals gene family evolution of the group 1 allergens in astigmatic mites. Clin. Transl. Allergy, 13, e12324.

Trachana, K. et al. (2011) Orthology prediction methods: a quality assessment using curated protein families. BioEssays, 33, 769–780.

Vizueta, J. et al. (2020) BITACORA: a comprehensive tool for the identification and annotation of gene families in genome assemblies. Mol. Ecol. Resour., 20, 1445–1452.

Wang, J. et al. (2023) The conserved domain database in 2023. Nucleic Acids Res., 51, D384–D388.

