## Supplementary Information for "GeneFamHunter: annotation-independent recovery of divergent and tandemly arrayed gene-family members from genome assemblies"

Contents: Note S1, fused gene models in reference proteomes (Table S1, Fig. S1). Note S2, window-based searches in six QfO genomes (Tables S2–S3, Figs S2–S4). Note S3, window-based searches in six mite genomes (Tables S4–S5, Figs S5–S7). Note S4, full-pipeline comparison with Shi et al. (2023) and the authors' automated annotations, and the MEROPS-labelled gene tree (Tables S6–S7, Fig. S8). Note S5, clustering parameters (Table S8). Note S6, family membership compared with orthology groups (Tables S9–S10, Fig. S9).

### Supplementary Note S1. Artificially fused gene models in reference _proteomes_

#### Methods

Here, we focus on cysteine protease genes, but this strategy can be applied to any gene family in which each gene contains no more than one copy of the target domain, even if the encoded proteins contain other domains. Cysteine protease genes usually contain an inhibitor domain and a single catalytic domain, PF00112 (ref). However, these genes can be tandemly arrayed in the genome, and gene prediction software may misidentify a stop codon as an intron, artificially extending a gene model into an adjacent locus. As a result, a single predicted protein may contain multiple catalytic domains, although published studies indicate that cysteine protease genes typically encode a single PF00112 catalytic domain (ref). Such artificially fused genes are present in reference proteomes and may propagate annotation errors in downstream analyses.

To evaluate the extent of these fused proteins, we screened reference proteomes using HMMER v3.3.2 and the PF00112 profile as part of the GeneFamHunter preparation step (run_prep_HMM.sh, function find_multidomain_prots; main text Fig. 1A, step 7). A protein was flagged when it contained two or more PF00112 regions separated by more than 20 residues. The protein evidence used for gene prediction, designated uniprot.5spp, comprised the curated UniProt Swiss-Prot database and five mite proteomes derived from chromosome-level genome assemblies: *Blomia tropicalis* (GCA_021650745.2), *Tyrophagus putrescentiae* (GCA_021730765.1), *Sarcoptes scabiei* (GCA_020844145.1), *Dermatophagoides farinae* (GCF_020809275.1), and *Dermatophagoides pteronyssinus* (GCF_001901225.1). The proteomes of the first three mite species were generated by researchers using gene-prediction tools such as MAKER, whereas those of the latter two were obtained from GenBank's annotation pipeline. Thus, the dataset combines a manually curated reference protein database with a small, targeted sample of mite proteomes produced through genome annotation.

Protein sequences were concatenated into a single dataset, retaining source-specific identifiers, and searched with hmmscan to identify PF00112 domains. Proteins containing a single PF00112 domain were classified as single-domain candidates, whereas those containing multiple copies were flagged as potentially fused models. Proteins with multiple PF00112 domains were excluded from the filtered reference dataset, and single-domain proteins were retained as a high-confidence query set for downstream analyses, including gene prediction with GeneFamHunter. To separate papain-like proteins (C1A) from bleomycin hydrolases (C1B), which PF00112 also detects, every PF00112 hit was also scored with PF03051 (Peptidase_C1_2; hmmsearch, HMMER 3.3.2), and proteins with a higher full-sequence bit score for PF03051 were classified as C1B. The C1B proteins were kept in the gene-prediction evidence.

This filtering strategy is intended only for gene families in which a single catalytic domain is expected per gene. It should not be applied to multidomain proteins in which repeated homologous domains represent the native biological architecture, such as pseudo-dimeric proteins whose duplicated domains fold together to form a single functional active site.

#### Results

Across the 645,201 proteins in our concatenated dataset (uniprot.5spp), PF00112 detected 466 proteins, 190 from the five mite proteomes and 276 from Swiss-Prot. Twenty Swiss-Prot proteins scored higher with PF03051 and were classified as C1B (bleomycin hydrolases, yeast BLH1 and bacterial aminopeptidases C, E, G and W), leaving 256 Swiss-Prot and 190 mite C1A proteins. In the GenBank sequences of the five mite species, 18 of the 190 C1A proteins were putative fused genes with more than one PF00112 domain (9.47%), whereas none of the 256 Swiss-Prot C1A proteins was fused (Table S1). Of the 18 fused GenBank gene models, 14 were generated by MAKER annotation, while four were produced by the NCBI Eukaryotic Genome Annotation Pipeline (Supplementary Data X1). Fig. S1 illustrates a representative example (XP_046918738.1). At the boundary between the two underlying genes, RNA-seq coverage drops, consistent with a gene boundary that was incorrectly annotated as an intron. This annotation joins two tandemly arranged genes into a single artificial chimeric gene model. Other examples include gene models containing seven (KAI2799262.1) or five (KAH9396860.1) artificially fused domains (Supplementary Data X1: protein identifiers; X2: protein sequences).

Overall, approximately 9.5% of C1A gene models in the five GenBank genomes appear to be artificially fused, indicating that such misannotations may propagate into downstream analyses. In contrast, the Swiss-Prot database did not contain fused cysteine protease gene models. To avoid predicting artificially fused genes, we removed the fused gene models from our protein evidence dataset prior to running the GeneFamHunter pipeline (see Methods).

We applied the same screen (hmmsearch with PF00112, -Z 1 --domZ 1, i-E-value < 0.01; find_multidomain_prots with a 20-residue gap) to the six Quest for Orthologs proteomes used in Note S2 (release 2026_02; 94,853 canonical proteins and 20,284 isoforms). Of 148 PF00112 hits, six were C1B (the bleomycin hydrolases X2JEJ0, D2A4S5 and A0A0P5X8K8 and three isoforms of X2JEJ0). Of the remaining 142 C1A proteins (the 137 UniProt IPR000668 proteins, four of their isoforms without the IPR000668 annotation, and Q8IBZ2, an α/β-hydrolase with a weak PF00112 hit), 2 (1.4%; 2 of the 128 canonical C1A proteins, 1.6%) were putatively fused: D6WR26 and A0A139WAQ9, both unreviewed T. castaneum entries carrying two tandem cathepsin L units. No fused proteins were found in the other five proteomes. Both proteins are part of the QfO reference, where each contributes two domain loci (Note S2).

Table S1. PF00112 proteins in the unfiltered reference proteome (uniprot.5spp) and, for comparison, in the six QfO proteomes. Proteins scoring higher with PF03051 were classified as bleomycin hydrolases (C1B); the remaining papain-like (C1A) proteins are split into single-domain and putative fused gene models (two or more PF00112 domains). Before gene prediction, proteins with multiple PF00112 domains are filtered out by the GeneFamHunter pipeline.

| **Subset** | **GenBank mites** | **UniProt Swiss-Prot** | **QfO proteomes (6 species)** |
| --- | --- | --- | --- |
| Proteomes concatenated* | 74,371 | 570,830 | 115,137 |
| PF00112 hits | 190 | 276 | 148 |
| Bleomycin hydrolases (C1B; PF03051 > PF00112) | 0 | 20 | 6 |
| Papain-like (C1A) | 190 | 256 | 142 |
| Single-domain C1A | 172 | 256 | 140 |
| Putative fused C1A | 18 | 0 | 2 |
| Fused C1A, % | 9.47 | 0.00 | 1.41 |

Protein evidence for gene prediction (uniprot.5spp) included the UniProt Swiss-Prot database and predicted mite proteomes from five mite species (with the 18 fused proteins removed): *Blomia tropicalis* (GCA_021650745.2), *Tyrophagus putrescentiae* (GCA_021730765.1), *Sarcoptes scabiei* (GCA_020844145.1), *Dermatophagoides farinae* (GCF_020809275.1), *Dermatophagoides pteronyssinus* (GCF_001901225.1). QfO proteomes: canonical proteins plus isoforms of the six genomes in Note S2; they were screened for comparison and are not part of the gene-prediction evidence

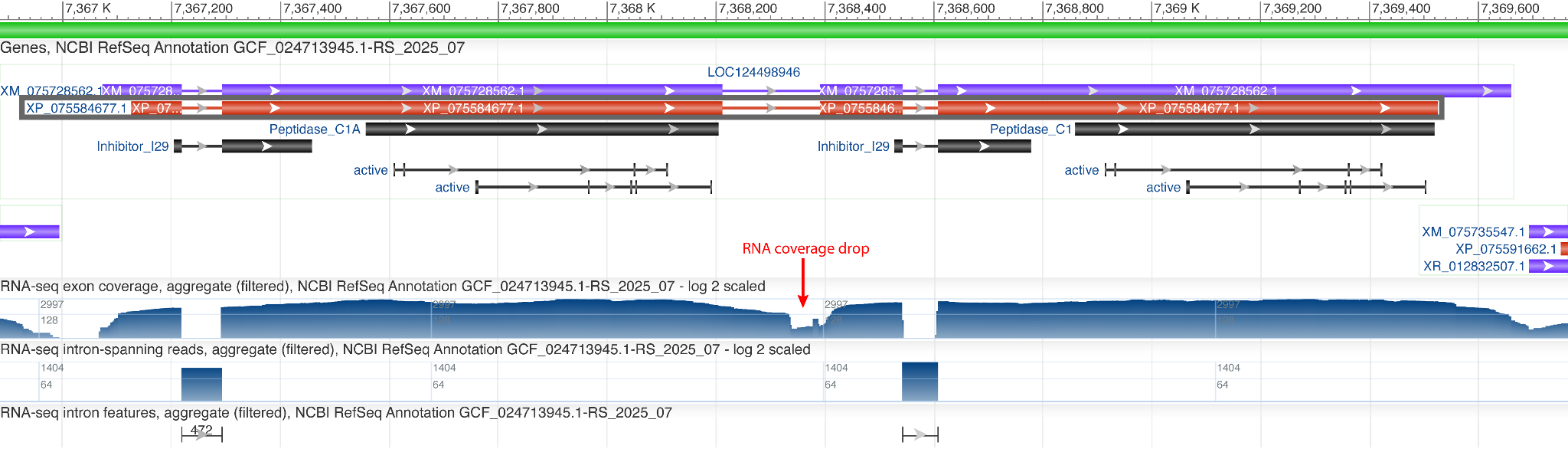

**Fig. S1.** re X1. An artificially fused gene model in the house dust mite, Dermatophagoides farinae, produced by the NCBI Eukaryotic Genome Annotation Pipeline. Each of the two underlying genes contains the expected Inhibitor_I29 and cysteine protease Peptidase_C1 domains. A drop in RNA-seq coverage near the boundary between the underlying genes (red arrow) is consistent with a gene boundary that was incorrectly annotated as an intron, thereby joining the two genes into a single predicted model (LOC124498946). This model corresponds to the fused protein XP_046918738.1, using the earlier annotation for compatibility with the published study (Shi et al., 2023). The fused model was also retained in the current (2025) release of the NCBI GenBank annotation (shown here).

### Supplementary Note S2. Window-based searches in six Quest for Orthologs genomes

#### Dataset

Our Quest for Orthologs (QfO) dataset includes six species broadly distributed across the eukaryotic tree of life: Plasmodium falciparum (accession), Dictyostelium discoideum, Caenorhabditis elegans, Drosophila melanogaster, Tribolium castaneum, and Daphnia magna.

To compare TBLASTN with the other homology-search methods under conditions where closely related protein queries are unavailable, we used distantly related mite group 1 allergen proteins as queries. This setup represents a challenging but realistic scenario in which only distant homologs of the target gene family are available. For reproducibility, the genomes and queries are documented as part of our test scripts (GeneHunter_benchmarks repository).

#### Methods

We evaluated how genomic target construction affects the sensitivity and specificity of HMMER homology searches, focusing on the recovery of conserved protein domains from genomic assemblies while minimizing additional hits outside known loci. Specifically, we tested whether six-frame continuous translation of overlapping genomic windows, as implemented in GeneFamHunter, can recover domains spanning introns containing in-frame stop codons, which potentially fragment translated targets in ORF-based approaches such as TGFam-Finder. We also examined how window size affects domain recovery and merged-hit length, particularly for short terminal exons, and how HMMER acceleration filters (--max) affect the recovery of weakly scoring domain fragments.

We compared five HMMER search strategies: (E1) six-frame translation of 1,400-bp sliding windows with 700-bp overlap (BEDTools 2.31.0, makewindows -w 1400 -s 700; SeqKit 2.3.1, translate -f 6), retaining stop codons as * to produce continuous translations. This strategy is implemented in GeneFamHunter, where window size and overlap can be configured according to the expected target size; (E2) identification of stop-to-stop open reading frames (ORFs) in the six-frame translation of the genomic assemblies, retaining only ORFs of at least 20 codons. Each ORF is bounded by stop codons and treated as a separate protein-like target for HMMER searches; (E3) the same ORF-based search with all HMMER acceleration filters disabled (--max); (E4) six-frame translation of 290-kb windows overlapping by 10 kb; and (E5) the same long-window search with all acceleration filters disabled (--max). The long-window strategy provided a comparison with substantially longer genomic targets, given HMMER target-sequence length limit of approximately 100,000 residues for the searches used here. All five strategies used the same 220-position PF00112 cysteine protease catalytic-domain profile, identical search settings (-Z 1 --domZ 1) fixing the effective target-database size for compatible E-value calculations, and a domain inclusion threshold of i-Evalue < 0.01. Before analysis, we excluded 17 Tribolium castaneum ORFs exceeding the target-length limit; all consisted of at least 99.96% X characters, representing assembly gaps lacking stop codons.

The primary reference (truth I) comprised 130 papain-like (C1A) domain loci (InterPro IPR000668, the family modelled by PF00112; 128 canonical proteins and nine isoforms, with the two fused proteins contributing two loci each; bleomycin hydrolases, C1B, are not included) in six eukaryotic genomes from the Quest for Orthologs (QfO) dataset. The loci were identified using official UniProt annotations and mapped to their corresponding genomic coding sequences, with protein isoforms collapsed. Because UniProt domain annotations are largely derived from InterPro analyses, which use HMMER, this reference may not be fully independent of profile-based searches. We therefore also evaluated recovery against a secondary reference (truth B) constructed independently of GeneFamHunter own HMMER search results. This reference comprised 133 regions of interest (ROIs) generated by conventional RPS-BLAST searches of whole contigs, alongside 21 unsupported ROIs retained for the broader assessment. This complementary reference allowed us to assess whether the relative performance of the search strategies held when evaluated against regions supported by a different search approach. The construction and evaluation of truth B are described in the GeneHunter_benchmarks repository (analyses/00_qfo_reference).

A reference locus was considered recovered when a merged genomic hit overlapped its annotated domain region by at least 30 bp on the same strand. For each search, hits on the same contig and strand that overlapped in genomic coordinates were joined into merged hits. We counted the number of reference loci recovered and the number of merged hits outside all reference loci; the latter may include partial gene hits and were not assumed to be false positives.

For E1 and E2, we additionally compared the genomic positions of merged hits recovered by only one strategy and mapped their alignments to the 220-position PF00112 model. For merged hits shared by both strategies, we assessed whether each E1 hit corresponded to a single E2 hit or spanned two E2 hits separated by a gap. In the latter cases, we inspected the translated genomic windows to determine how the window-based alignment crossed the intervening region. Finally, among merged hits within reference loci, we compared genomic hit length and PF00112 model coverage (hmm_to - hmm_from + 1, expressed relative to the 220-position model) between each strategy and E1 using two-sided Wilcoxon rank-sum tests.

We additionally compared RPS-BLAST, PSI-BLAST, DELTA-BLAST, and TBLASTN searches using 1,400-bp windows and whole contigs. TBLASTN was included for comparison as it is often used in gene recovery pipelines, but is not part of GeneFamHunter. RPS-BLAST used the pfam00112 CDD profile. To assess performance when closely related query proteins are unavailable, PSI-BLAST and DELTA-BLAST were run using the 11 mite group 1 allergen proteins as queries against the six test genomes.

The code documenting these analyses and datasets has been deposited in tests/windows_vs_contigs_qfo.

#### Results

We evaluated how genomic target construction affects the sensitivity and specificity of homology searches: six-frame continuous translation of overlapping 1400-bp genomic windows (E1), ORFs, interrupted by stop codons (E2), and large, 290-kb windows (E4), resembling a whole contig search approach. For E2 and E4, we ran two additional analyses with HMMER acceleration filters disabled (E3 and E5, respectively). Four of the five searches (E1–E3 and E5) recovered all reference loci, whereas the 290-kb window search (E4) recovered only 83.1% (main text Fig. 1C; Table S2). Among the four searches that recovered all loci, the 1,400-bp window approach (E1) produced the fewest merged hits outside reference loci, followed by the ORF-based approach (E2), with 21 and 45 such hits, respectively (main text Fig. 1C; Fig. S2B; Table S2).

We further compared E1 and E2 to examine differences in the recovery of terminal exons. Within reference loci, E1 yielded 10 hits not recovered by E2, with relatively weak i-E-values (3 × 10⁻⁶ to 5 × 10⁻³), whereas E2 yielded five hits not recovered by E1, with stronger i-E-values (2 × 10⁻¹² to 7 × 10⁻⁶). The E1-specific fragments were concentrated near the ends of the HMM model coordinates, consistent with their representing terminal exons (Fig. S2A: blue). In E1, these fragments are translated together with the main domain, potentially allowing the combined sequence to pass HMMER per-sequence filters. In contrast, E2-specific hits were distributed more evenly across the HMM model coordinates (Fig. S2A: green). Disabling HMMER acceleration filters for the ORF search (E3) recovered 10 of the 12 E1-specific merged hits across both reference loci and regions outside them. These results suggest that the difference between E1 and E2 partly reflects the effects of HMMER acceleration filters on the detection of weak domain fragments in terminal exons. Disabling the filters (E3) substantially increased recovery but also produced a very large number of additional hits outside reference loci (Table S2).

Hit lengths were similar between window- and ORF-based searches (Fig. S3). Within reference loci, E1, E2, and E3 produced similar median hit lengths (225–239 bp) and median model coverage (30–31%), with no significant differences from E1 in either measure (Wilcoxon rank-sum tests, P ≥ 0.11). E5 (290-kb windows with acceleration filters disabled) also did not differ significantly from E1. In contrast, E4 (290-kb windows with default filters) produced longer merged hits (median 278 bp; 35% model coverage; P = 0.031 and 0.032, respectively), partly because this search failed to recover short, weakly scoring fragments.

Of the 297 E1 merged hits that overlapped E2 hits, 17 hits each spanned two E2 merged hits separated by a short gap (median, 33 bp; range, 3–217 bp) containing an in-frame stop codon that an ORF-based search could not cross. In E1, the alignment spanned the stop codon(s) in 11 cases; in the other six, the envelope of one alignment extended beyond a stop codon to the adjacent fragment. Thus, translating across stop codons allowed window-based searches to recover some interrupted domains as single hits, but did not result in systematically larger merged hits (Fig. S3).

We further investigated the 290-kb window search (E4), which failed to recover 22 reference loci (Table S2). The missed loci occurred in translated targets averaging approximately 75,000 residues per frame in D. discoideum and 85,000 in P. falciparum. Disabling HMMER acceleration filters (E5) restored recovery of all reference loci. In a separate comparison of 227 merged hits paired one-to-one between the 290-kb and 1,400-bp window searches, the median i-E-value was approximately 10²·⁵-fold higher for the longer windows (Fig. S4B).

In addition to the HMMER benchmarks, we compared searches of 1,400-bp sliding windows and whole contigs using four NCBI tools representing domain-, profile-, and similarity-based approaches: RPS-BLAST, PSI-BLAST, DELTA-BLAST, and TBLASTN (main text Fig. 1D; Table S3; Fig. S4). We evaluated reference-locus recovery, merged hits unique to each target representation, and differences in E-values.

BLAST-based tools generally recovered the same loci with or without windowing (Table S3; Fig. S4A). But the recall of these methods was different. PSI-BLAST and DELTA-BLAST each recovered all 130 reference loci with both target representations, producing nearly identical sets of merged hits and no substantial shift in E-values (Fig. S4B). RPS-BLAST showed a more pronounced difference: window-based searches recovered 129 loci (99.2%), compared with 128 (98.5%) for whole-contig searches (Table S3). For shared hits, the median E-value was approximately 10^4.4-fold higher in whole-contig searches than in window-based searches (Fig. S4B). Thus, windowing had only a modest effect on locus recovery, while substantially improving the reported E-values and increasing the number of detected hits. TBLASTN recovered 123 loci with windows (94.6%) and 124 (95.4%) with whole contigs (Table S3). We therefore used the more sensitive PSI-BLAST rather than TBLASTN in GeneFamHunter.

Conclusion. Across six QfO genomes, using a reference independent of GeneFamHunter search results, HMMER searches of 1,400-bp sliding windows recovered all reference domain loci while producing the fewest merged hits outside them. ORF-based searches also recovered all loci but produced more than twice as many hits outside the reference loci and missed some short domain fragments, including terminal-exon fragments. While whole-contig searches are not possible because of the HMMER target-length constraint, our longest tested windows (290 kb) recovered only 83.1% of reference loci with default acceleration filters. In contrast, windowing had little effect on reference-locus recovery by the BLAST-based tools, except that windowed rpstblastn searches yielded additional hits, consistent with the effects of applying thresholds to shorter targets. These results support GeneFamHunter use of configurable, overlapping genomic windows for HMMER searches and provide a consistent translated-target representation across search tools for general homology sensitivity search applications.

Table S2. HMMER searches scored against truth I (UniProt annotations, 130 IPR000668 domain loci in the six QfO genomes) and, in brackets, against truth B (conventional rpstblastn search of whole contigs, 133 ROIs).

| **#** | **Search** | **Merged hits** | **Recall, truth I (truth B)** | **Recall %, truth I (truth B)** | **Merged hits outside reference loci** |
| --- | --- | --- | --- | --- | --- |
| E1 | 1,400-bp windows | 309 | 130 (133) | 100.0% (100.0%) | 21 |
| E2 | ORFs | 338 | 130 (133) | 100.0% (100.0%) | 45 |
| E3 | ORFs, filters off | 108,124 | 130 (133) | 100.0% (100.0%) | 107,801 |
| E4 | 290-kb windows | 210 | 108 (110) | 83.1% (82.7%) | 8 |
| E5 | 290-kb windows, filters off | 583 | 130 (133) | 100.0% (100.0%) | 317 |

Table S3. BLAST-based tools on 1,400-bp and on whole contigs, with the same commands and parameters as in Table S2, scored against truth I (truth B in brackets). For rpstblastn, truth B is defined by the whole-contig run itself. For protein queries (psiblast, deltablast and tblastn) we used 11 distant homologs (mite group-1 allergens), so recall for these tools is not comparable with that of the profile-based tools (rpstblastn). tblastn is not part of GeneFamHunter.

| **Tool** | **Input** | **Merged hits** | **Recall, truth I (truth B)** | **Recall %, truth I (truth B)** | **Merged hits outside reference loci** |
| --- | --- | --- | --- | --- | --- |
| rpstblastn | 1,400-bp windows | 208 | 129 (133) | 99.2% (100.0%) | 8 |
| rpstblastn | whole contigs | 189 | 128 (133) | 98.5% (100.0%) | 7 |
| psiblast | 1,400-bp windows | 232 | 130 (133) | 100.0% (100.0%) | 16 |
| psiblast | whole contigs | 228 | 130 (133) | 100.0% (100.0%) | 15 |
| deltablast | 1,400-bp windows | 279 | 130 (133) | 100.0% (100.0%) | 31 |
| deltablast | whole contigs | 278 | 130 (133) | 100.0% (100.0%) | 32 |
| tblastn | 1,400-bp windows | 197 | 123 (129) | 94.6% (97.0%) | 7 |
| tblastn | whole contigs | 203 | 124 (130) | 95.4% (97.7%) | 8 |

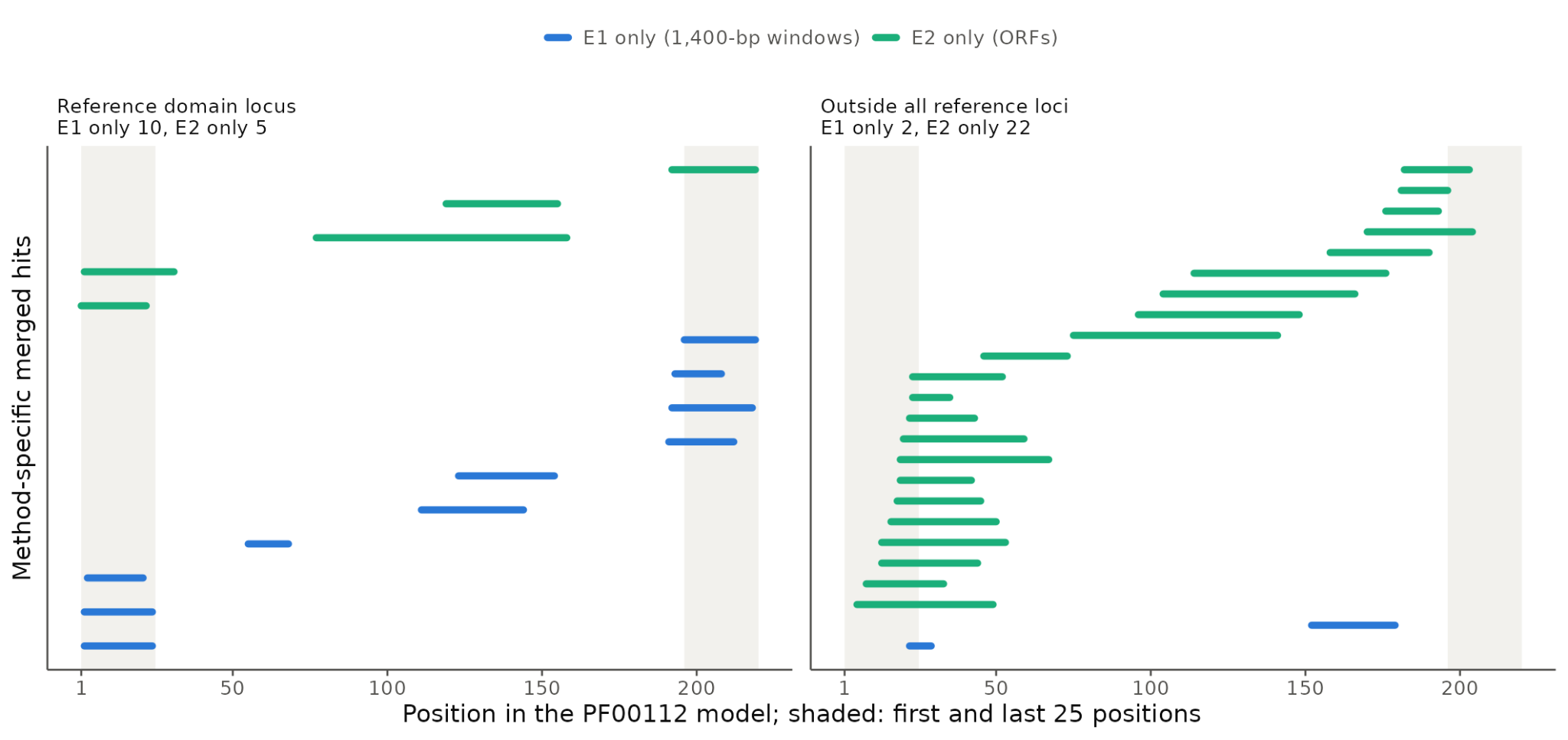

**Fig. S2.** Distribution of merged hits detected exclusively by E1 (1,400-bp windows) or E2 (ORFs) along the PF00112 model. Each line represents one merged hit. Blue, E1-specific hits; green, E2-specific hits. Panels show merged hits within reference domain loci (A) and outside all reference loci (B). Shading indicates the first and last 25 positions of the model.

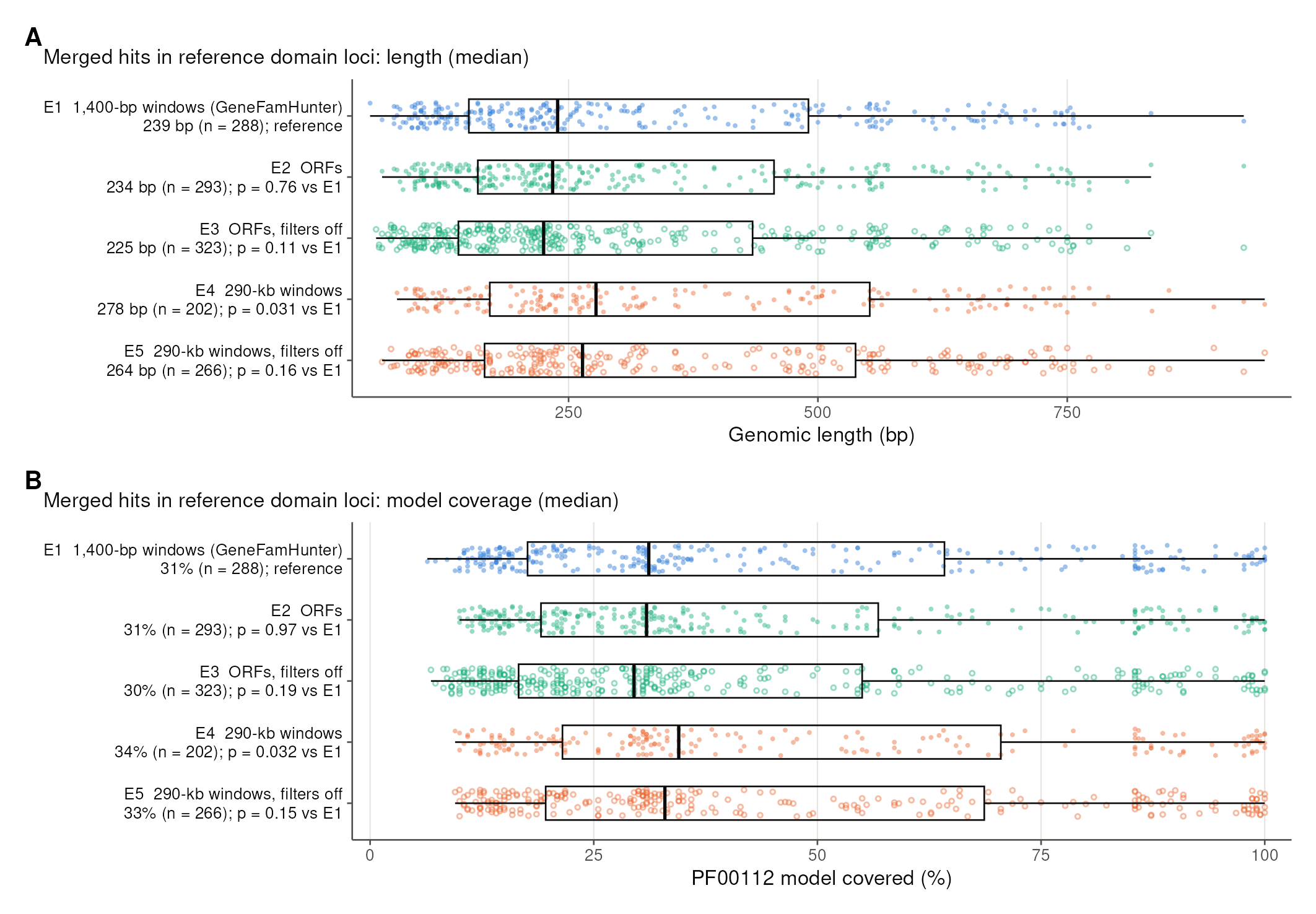

**Fig. S3.** Size of merged hits. (A) Genomic length and (B) PF00112 model coverage of merged hits inside reference loci, all genomes pooled. Axis labels give the median, n, and the two-sided Wilcoxon rank-sum p-value against E1. Boxes show the median and interquartile range; whiskers extend to 1.5 × IQR; open symbols mark searches with filters off.

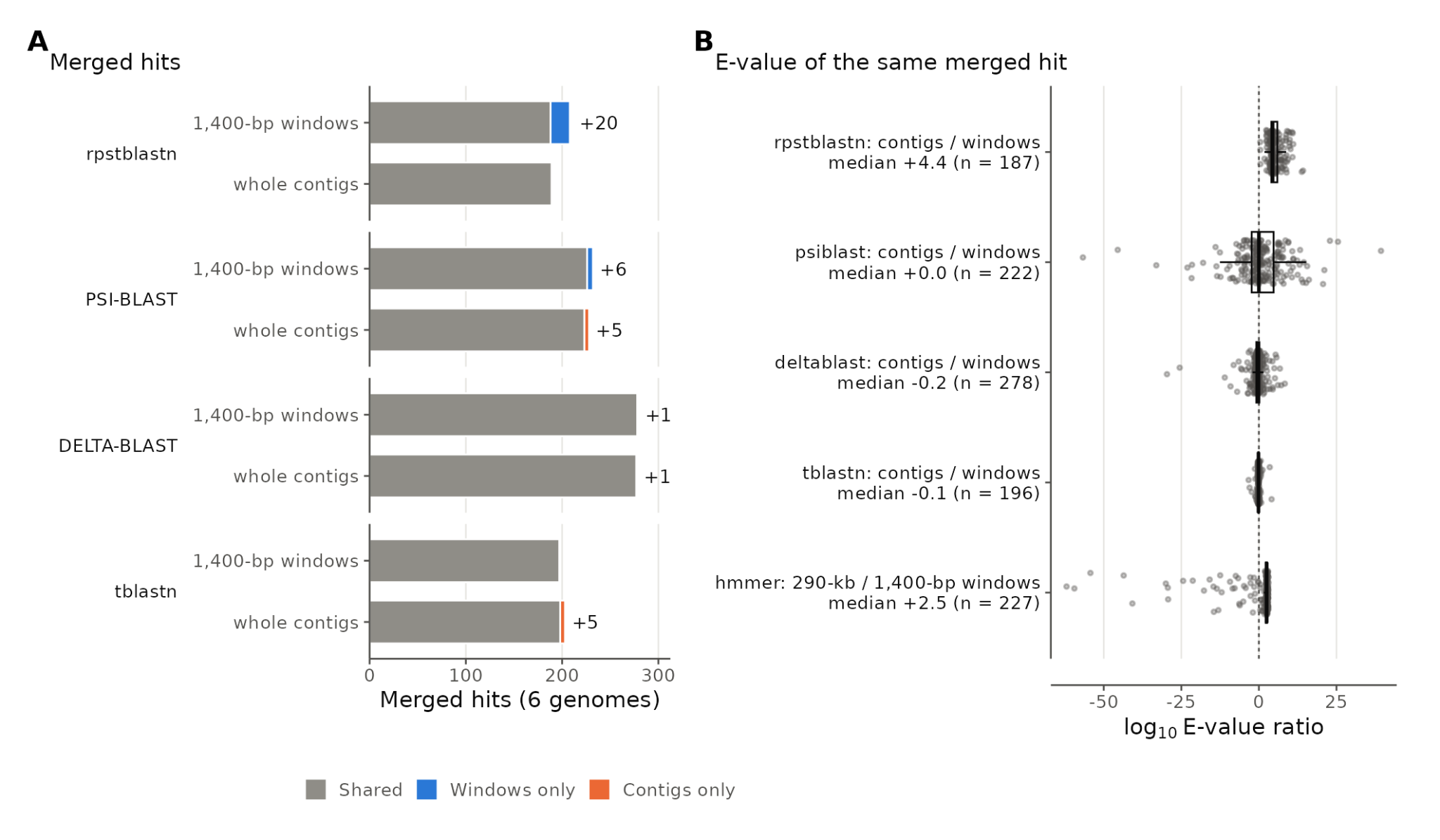

**Fig. S4.** ure 5. BLAST-based tools on 1,400-bp windows and on whole contigs. (A) Merged hits found on both inputs (grey), only on windows (blue) or only on contigs (orange). psiblast, deltablast and tblastn use the same 11 allergen queries. tblastn is not part of GeneFamHunter, but included here for comparison; (B) For merged hits paired one to one between the two inputs, log₁₀ of the E-value ratio (alternative / windows). The HMMER comparison of 290-kb with 1,400-bp windows is shown for reference. A positive value means the alternative scores the same hit worse.

### Supplementary Note S3. Window-based searches in six mite genomes

#### Methods

This benchmark extended the above QfO window-versus-whole-contig analysis (Note S2) to six species of mites causing allergies (Shi et al., 2023). Because these genomes lack an independent annotation of the cysteine protease C1 family, searches were evaluated against regions of interest (ROIs) identified by GeneFamHunter and supported by rpstblastn (truths A and B; Table S4). We did not use GeneMark-predicted gene models as the evaluation reference because variation in gene-prediction accuracy could confound the assessment of candidate-region detection by introducing errors attributable to gene-model reconstruction. The ROI-based reference is not fully independent of GeneFamHunter and may favor its search strategy. The same scripts were used as in the primary benchmark.

#### Results

We extended the QfO benchmark to six allergy-inducing mite species (Shi et al., 2023). Because independent annotations of the cysteine protease C1 family were unavailable, evaluation used GeneFamHunter-derived regions of interest (ROIs) supported by rpstblastn searches of whole contigs (truth B; 246 true-positive and 23 false-positive ROIs) or of 1,400-bp windows (truth A; 253 and 16 ROIs). Truth B is the primary reference; truth A is given in brackets. Because E1 is part of the GeneFamHunter pipeline, its absence of hits outside the ROIs is guaranteed by construction.

The results showed a pattern consistent with the primary QfO benchmark: E1 (1,400-bp windows), E2 (ORFs), and both searches with HMMER acceleration filters disabled (E3 and E5) recovered all true-positive ROIs, whereas E4 (290-kb windows with default filters) did not. E4 recovered a larger proportion of true-positive ROIs in the mite genomes (90.2% under truth B and 89.7% under truth A) than in the QfO genomes (83.1% under truth I and 82.7% under truth B) (Table S4). In both benchmarks, median i-E-values were lower for short windows than for long windows (Figs S4B and S7), indicating that short-window searches yielded more statistically significant matches and supporting their use over long windows. Disabling the acceleration filters restored complete recovery in both benchmarks. In other words, HMMER cannot search unwindowed genomes at all, and with default filters it loses real domains in the longest targets it accepts.

In terms of terminal-exons, as in the QfO benchmark, E1 and E2 differed in their recovery of terminal-exon fragments, with a more pronounced effect in the mite benchmark. E1-only fragments were generally short, weak, and concentrated near the ends of the HMM model (Fig. S5A: blue), consistent with terminal exons. When searched as separate ORFs, these fragments may fail to pass HMMER acceleration filters (Fig. S5A: green). Disabling the filters for the ORF search (E3) recovered all but one of the 25 E1-only merged hits, supporting the interpretation that HMMER filtering contributes to the difference between window- and ORF-based searches.

In terms of median hit length, the mite benchmark showed a pattern broadly consistent with the QfO benchmark (Figs S3 and S6). In both datasets, median hit length and model coverage were lower for E2 than E1 and lower still for E3, whereas both long-window searches (E4 and E5) yielded higher medians. Disabling HMMER acceleration filters reduced median hit length and model coverage relative to the corresponding filtered searches (Fig. S6). Although absolute medians and statistical significance relative to E1 differed between datasets, the overall patterns were consistent. Translating across stop codons allowed window-based searches to recover some interrupted domains as single hits, yielding numerically greater median hit lengths and model coverage than ORF-based searches, although neither difference was statistically significant (E1: P = 0.087 and E2: P = 0.24).

The BLAST-based tools generally recovered the same true-positive ROIs with 1,400-bp windows and whole-contig targets (Table S5; Fig. S7). PSI-BLAST and DELTA-BLAST achieved 100% recall with both target representations under both truths. rpstblastn also recovered all ROIs except that, under truth A (defined by the windows run), the whole-contig search missed seven window-defined ROIs (246 of 253; 97.2%); each truth favours the rpstblastn run that defines it. TBLASTN showed lower recall, recovering 233 of 246 true-positive ROIs (94.7%) with windows and 232 of 246 (94.3%) with whole contigs. This lower recall likely reflects the limited representation of the C1 family by the 11 mite group 1 allergen queries, rather than an effect of windowing. Unlike these query-dependent searches, rpstblastn uses conserved-domain profiles to detect homologous regions. Although recall was largely unchanged, target representation affected the number of merged hits. rpstblastn produced 26 window-only merged hits, but these did not identify any additional true-positive ROIs beyond those recovered from whole contigs. This difference is consistent with the more permissive, window-local E-value threshold, which makes some alignments more likely to pass the significance cutoff when searched against short windows. PSI-BLAST produced more contig-only than window-only hits (11 versus 2), while DELTA-BLAST showed little difference (2 contig-only hits and none unique to windows). TBLASTN results were also similar between target representations.

As in the QfO benchmark, based on E-values, RPS-BLAST favored window-based searches, with the median E-value approximately 10^3.9-fold higher (Fig. S7). There were no pronounced differences in E-values for DELTA- and TBLASTN searches, while e-values for PSI-BLAST favored contig search (Fig. S7).

Overall, windowing did not improve BLAST-based recall in this benchmark. Its principal justification lies instead in accommodating HMMER target-length constraints and providing a consistent translated-window representation across search methods.

Conclusion. In six non-model mite genomes, HMMER searches of 1,400-bp sliding windows, ORF-based searches, and searches of long windows with acceleration filters disabled recovered all reference ROIs, whereas 290-kb windows with default filters recovered approximately 90%. As in the QfO benchmark, short-window searches yielded more statistically significant matches than long-window searches, and disabling acceleration filters restored complete ROI recovery. Window-based searches also recovered short domain fragments consistent with terminal exons that were sometimes missed by ORF-based searches. Median hit length and model coverage showed patterns broadly consistent with those in the QfO benchmark, with lower medians for ORF-based searches and higher medians for long-window searches.

BLAST-based tools generally recovered the same true-positive ROIs with 1,400-bp windows and whole-contig targets. Windowed rpstblastn searches produced additional merged hits and more statistically significant matches, but these additional hits did not increase ROI-level recall. PSI-BLAST, DELTA-BLAST, and TBLASTN showed little difference in ROI recovery between target representations.

Together, the two benchmarks support the use of configurable, overlapping genomic windows for HMMER searches, both to accommodate target-length constraints and to improve recovery of domain fragments, while providing a consistent translated-target representation across search methods. The non-model mite benchmark further demonstrates the applicability of this strategy to genomes lacking independent, comprehensive annotations of the target gene family.

Table S4. HMMER searches scored against regular rpstblastn on whole contigs (truth B: 246 true-positive and 23 false-positive ROIs) and rpstblastn on 1,400-bp windows (truth A: 253 true-positive and 16 false-positive ROIs; in brackets). Unsupported merged hits outside ROIs and are the same under both truths. For E1, this category is guaranteed to be 0 by construction.

| **#** | **Search** | **Merged hits** | **TP ROIs** | **Recall, truth B (truth A)** | **FP ROIs hits, truth B (truth A)** | **Merged hits outside ROIs** |
| --- | --- | --- | --- | --- | --- | --- |
| E1 | 1,400-bp windows | 437 | 246 (253) | 100.0% (100.0%) | 23 (16) | 0 |
| E2 | ORFs | 475 | 246 (253) | 100.0% (100.0%) | 21 (14) | 24 |
| E3 | ORFs, filters off | 93,887 | 246 (253) | 100.0% (100.0%) | 23 (16) | 93,333 |
| E4 | 290-kb windows | 305 | 222 (227) | 90.2% (89.7%) | 10 (5) | 0 |
| E5 | 290-kb windows, filters off | 406 | 246 (253) | 100.0% (100.0%) | 21 (14) | 8 |

Table S5. BLAST-based tools on 1,400-bp and on whole contigs, with the same commands and parameters as in Table S4. For rpstblastn, truth B is defined by the whole-contig run and truth A by the windows run. For protein queries (psiblast, deltablast and tblastn) we used 11 distant homologs (mite group 1 allergens), so recall for these tools is not comparable with that of the profile-based tools (rpstblastn). tblastn is not part of GeneFamHunter.

| **Tool** | **Input** | **Merged hits** | **Shared** | **Only this input** | **TP ROIs found, truth B (truth A)** | **Recall %, truth B (truth A)** | **FP ROIs hit, truth B (truth A)** | **Unsupported merged hits** |
| --- | --- | --- | --- | --- | --- | --- | --- | --- |
| rpstblastn | 1,400-bp windows | 337 | 311 | 26 | 246 (253) | 100.0% (100.0%) | 7 (0) | 0 |
| rpstblastn | whole contigs | 311 | 311 | 0 | 246 (246) | 100.0% (97.2%) | 0 (0) | 0 |
| psiblast | 1,400-bp windows | 330 | 328 | 2 | 246 (253) | 100.0% (100.0%) | 14 (7) | 0 |
| psiblast | whole contigs | 353 | 342 | 11 | 246 (253) | 100.0% (100.0%) | 20 (13) | 0 |
| deltablast | 1,400-bp windows | 395 | 395 | 0 | 246 (253) | 100.0% (100.0%) | 16 (9) | 0 |
| deltablast | whole contigs | 396 | 394 | 2 | 246 (253) | 100.0% (100.0%) | 16 (9) | 0 |
| tblastn | 1,400-bp windows | 356 | 352 | 4 | 233 (235) | 94.7% (92.9%) | 5 (3) | 0 |
| tblastn | whole contigs | 358 | 352 | 6 | 232 (234) | 94.3% (92.5%) | 6 (4) | 0 |

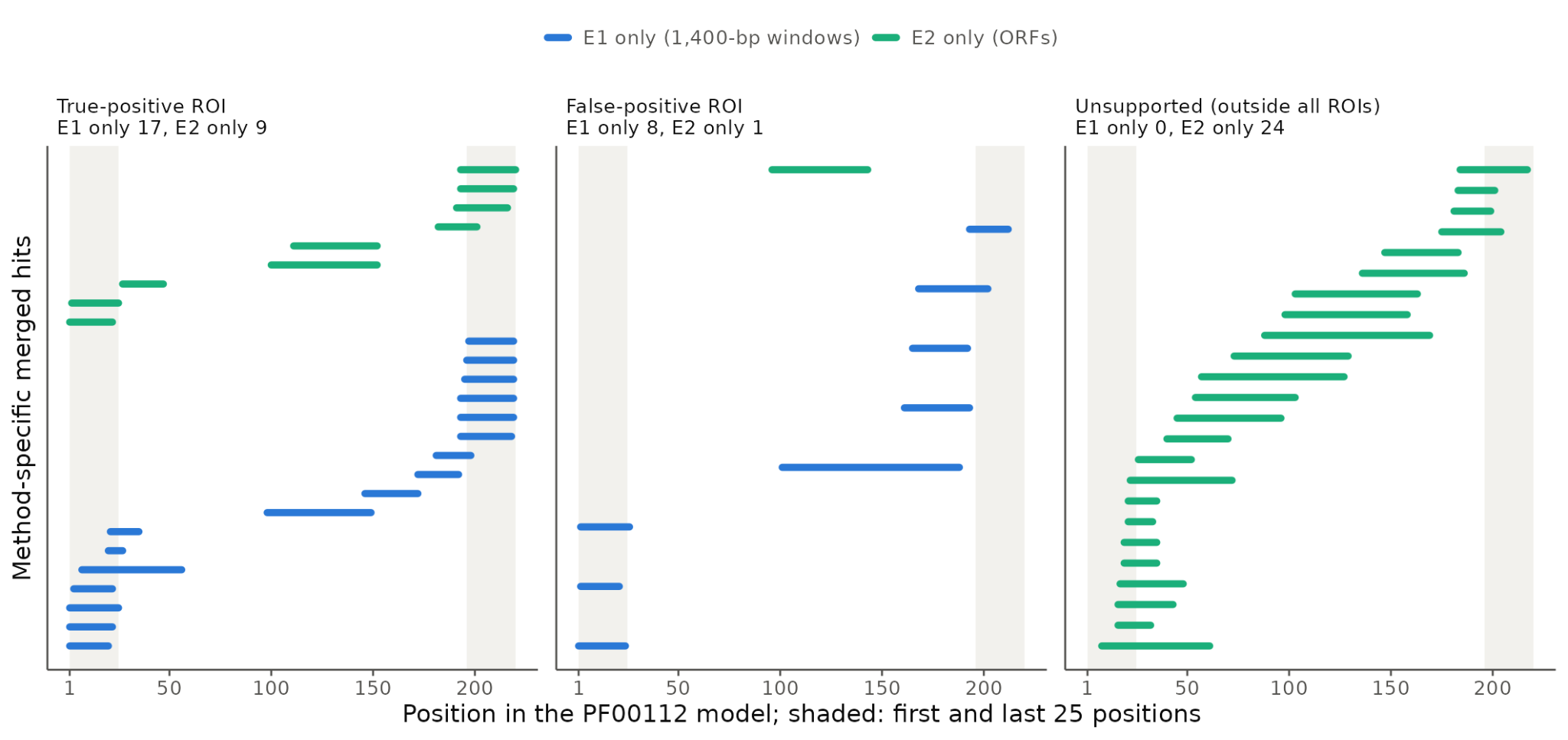

**Fig. S5.** re S2. Distribution of merged hits detected exclusively by E1 (1,400-bp windows) or E2 (ORFs) along the PF00112 model. Each line is one merged hit. Blue, E1-specific hits; green, E2-specific hits. Panels group merged hits by truth B: inside a true-positive ROI (A); inside a false-positive ROI (B); or outside all ROIs (C). Shading indicates the first and last 25 positions of the model.

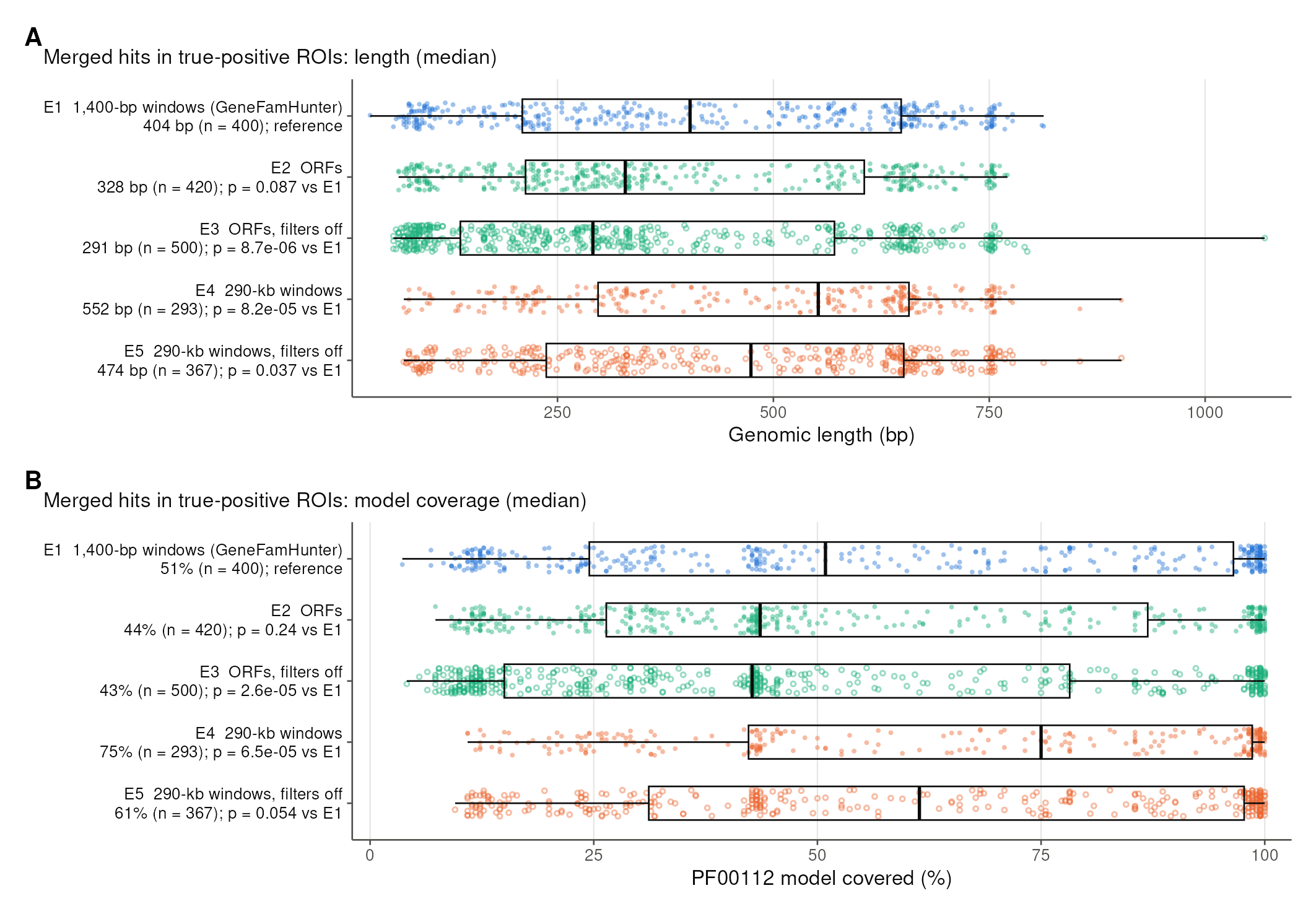

**Fig. S6.** Size of merged hits. (A) Genomic length and (B) PF00112 model coverage of merged hits inside true-positive ROIs (truth B), all genomes pooled. Axis labels give the median, n, and the two-sided Wilcoxon rank-sum p-value against E1. Boxes show the median and interquartile range; whiskers extend to 1.5 × IQR; open symbols mark searches with filters off.

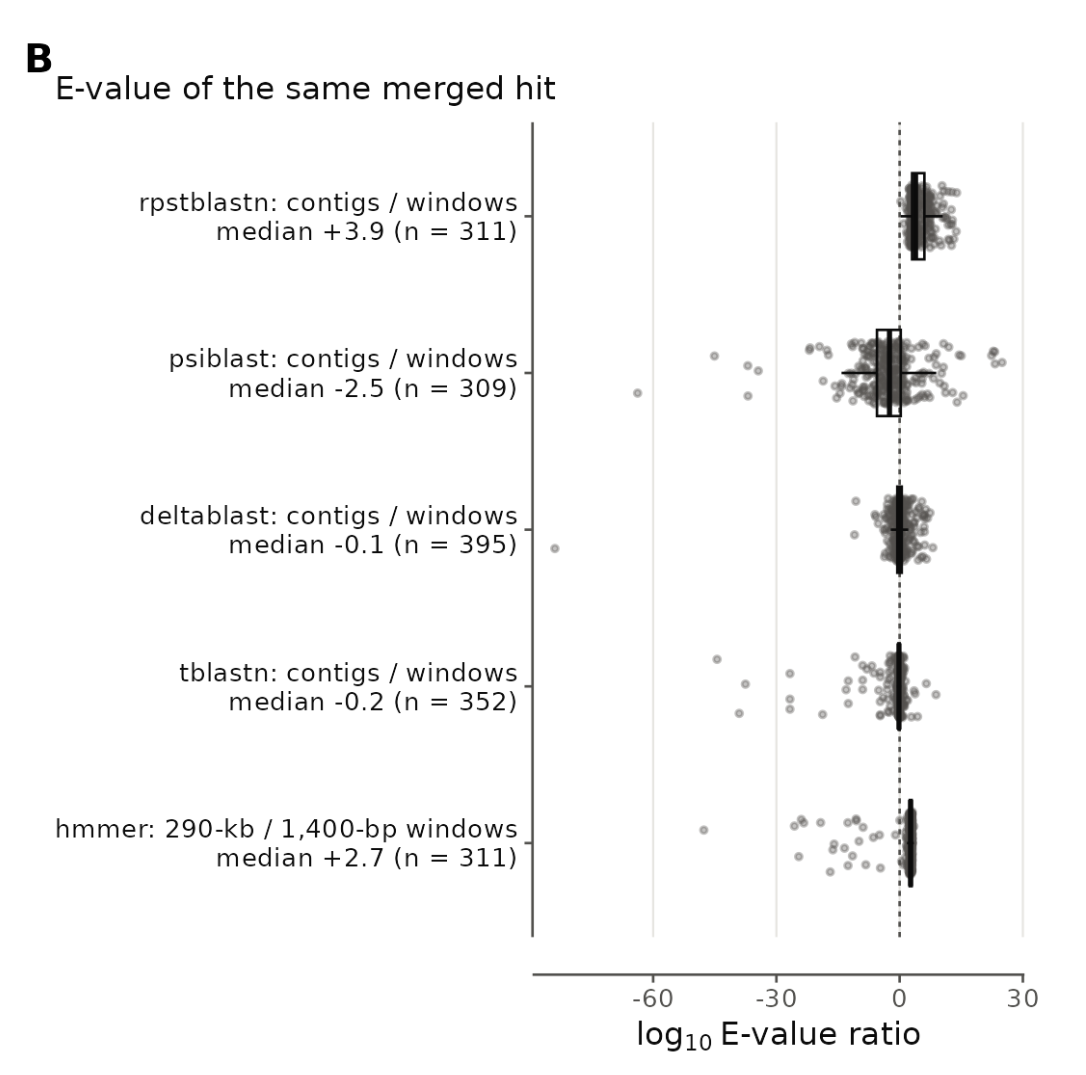

**Fig. S7.** BLAST-based tools on 1,400-bp windows and on whole contigs in six mite genomes. For merged hits paired one-to-one between the two inputs, log₁₀ of the E-value ratio (alternative / production windows). The HMMER comparison of 290-kb with 1,400-bp windows is shown for reference. A positive value means the alternative scores the same hit worse.

### Supplementary Note S4. Full-pipeline application to six mite genomes

#### Comparison with Shi et al. (2023)

The C1 set of Shi et al. (2023) comprises 217 proteins (163 gene identifiers; isoforms counted separately), which we placed on their genomes with the published GFF records or, where absent, with TBLASTN (E ≤ 10⁻⁵; HSPs merged when ≤ 3 kb apart and collinear in the query). Overlapping placements were collapsed into 211 genomic loci (B. tropicalis 50, D. farinae 26, D. pteronyssinus 28, P. ovis 22, S. scabiei 19, T. putrescentiae 66). A Shi locus was considered recovered when it overlapped a GeneFamHunter hit region (≥ 1 bp on the same contig) and, more strictly, a final GeneFamHunter gene model. Protein pairs were compared by BLASTP (E ≤ 10⁻²⁰): identical, ≥ 99% identity and ≥ 98% coverage of both proteins; minor difference, ≥ 95% identity and ≥ 90% coverage. Scripts: benchmarking1_OH_vs_Shi.v15/benchmarking2.sh.

| **Category (genomic hits, n = 271)** | **Loci** |
| --- | --- |
| Found by both (gene match 133; shared locus, model differs 75) | 208 |
| GeneFamHunter model below MIN_HIT_COV | 4 |
| GeneFamHunter region, no gene model | 18 |
| GeneFamHunter only | 39 (37 confirmed C1 genes; 12 full-length) |
| Shi23 non-C1 entries (no C1 domain in CDD) | 2 |

Table S6. Locus-level comparison between GeneFamHunter and the C1 set of Shi et al. (2023); this is a comparison, not a validation of either set. Two of the 211 Shi loci are Shi23 non-C1 entries: in a CDD search, DERF_002566 is a cysteine hydrolase family protein and TYRP_014064 has no conserved domain; neither has a GeneFamHunter hit. All 209 remaining loci overlapped GeneFamHunter hits, and 200 overlapped a final GeneFamHunter model (per species: BT 50/50, DF 25/25, DP 28/28, PO 22/22, SS 19/19, TP 56/65). GeneFamHunter-only loci per species: TP 22, BT 8, DP 6, DF 2, PO 1, SS 0; two of these (DP, PO) were classed as spurious_nt_hit and are not in the final set. At shared loci, protein models (n = 198) were identical (74), minor difference (54), GeneFamHunter truncated (29), divergent (34), Shi truncated (5) or Shi fused (2).

#### Comparison with the authors' automated gene annotations

PF00112 domains were identified with hmmsearch (HMMER 3.3.2; domain i-E-value < 0.01) in the automated gene annotations of the same six assemblies (99,699 proteins; funannotate for five species and MAKER for S. scabiei; provided by Shi et al. and not published or deposited in GenBank), and annotated C1 genes overlapping on the same strand were collapsed into loci. Each locus was matched to the 246 final GeneFamHunter models by genomic overlap on the same strand.

| **Species** | **Annotated C1 loci** | **GeneFamHunter models** | **Shared loci** | **GeneFamHunter only*** | **Annotation only** |
| --- | --- | --- | --- | --- | --- |
| B. tropicalis | 29 | 61 | 29 | 15 | 0 |
| D. farinae | 20 | 27 | 20 | 2 | 0 |
| D. pteronyssinus | 22 | 34 | 22 | 0 | 0 |
| P. ovis | 18 | 23 | 18 | 0 | 0 |
| S. scabiei | 6 | 20 | 6 | 14 | 0 |
| T. putrescentiae | 79 | 81 | 67 | 4 | 12† |
| Total | 174 | 246 | 162 | 35 | 12 |

Table S7. GeneFamHunter models versus C1 genes in the authors' automated annotations. *GeneFamHunter models that overlap no annotated gene of any kind. †Eleven of the 12 had GeneFamHunter evidence (ROIs) but were removed by the hit-coverage filter (3; 46.9–49.6% < 50%) or lacked a gene model (8). Twenty-eight annotated models contained 2–8 PF00112 domains and each spanned 2–8 separate GeneFamHunter models (76 in total; B. tropicalis 6 models → 23 genes, D. farinae 3 → 8, D. pteronyssinus 9 → 21, P. ovis 4 → 9, T. putrescentiae 6 → 15); one annotated model (BLOT_014423-RA) spanned eight. Such fused models were separated by manual curation in the published C1 set of Shi et al. (2023): only 2 of the 198 Shi proteins at shared loci remain fused (Table S6). When the GeneFamHunter proteins themselves were clustered with OrthoFinder 3.1.5 (default settings), 36 of 234 C1 proteins (15%) were not assigned to any orthogroup.

#### MEROPS-labelled gene tree

The 246 proteins were aligned with MAFFT and analysed with IQ-TREE 2 (VT+R8; 2,375 sites; 1,000 ultrafast bootstraps and SH-aLRT replicates). Each protein was assigned a MEROPS identifier by BLASTP against the MEROPS peptidase-unit library (release 12.5; E ≤ 10⁻⁵; high confidence, ≥ 50% identity and ≥ 70% unit coverage; medium, ≥ 35% identity), and labels were transferred to weakly matching tips through clades with SH-aLRT ≥ 80, UFBoot ≥ 95 and ≥ 80% label purity (merops_classify.sh). The identifiers are shown on the tips of the tree in Fig. S8 (merops_tree.R; R packages ape, treeio and ggtree). MEROPS classifies proteins only; tandem arrangement was read from the GeneFamHunter genomic coordinates, which are also part of each tip label.

The group 1 allergen homologs formed two supported clades: peptidase 1 (mite) (C01.073; clade C2: D. farinae 4, D. pteronyssinus 9, P. ovis 4, S. scabiei 10) and Blo t 1-type (C01.990; clade C3: B. tropicalis 4, T. putrescentiae 6). Many are tandemly arrayed: ten C01.073 copies within ~20 kb of S. scabiei scaffold3, four within ~11 kb of P. ovis PQWQ01000001.1, and four Blo t 1-type copies on T. putrescentiae SD3. Only four of these proteins match WHO/IUIS-listed allergens (blastp ≥ 95% identity; tagged [IUIS] in Fig. S8); the database lists clinically validated allergens only, so the remaining proteins are homologs, not demonstrated allergens. A supported clade of 16 T. putrescentiae genes (C18; SH-aLRT 98.4, UFBoot 99) had no confident MEROPS match; 14 of its members were found only by GeneFamHunter (marked *), and most are partial models. The tree agrees with the MEROPS classification despite many partial models (only 110 of 246 have both start and stop codons): 111 of the 114 proteins with their own MEROPS label that fall in supported clades share the clade's identifier, and 3 conflict. Seventeen proteins received their label from a supported clade; these are mostly partial models (3 of 17 complete; median 309 aa). The 51 proteins without any label are not shorter (median 401 aa; 10 complete) but lack close MEROPS homologs; 24 of them form the T. putrescentiae clades C18 (16 genes), C20 (5) and C21 (3). Clade C19 has no MEROPS hit of its own and contains exactly one gene from each of the six species (315–365 aa). It is the sister group of the cathepsin O genes within clade C4 (its tips carry the C4 label by clade transfer), so it is most likely a cathepsin O-related lineage without a close MEROPS homolog rather than a new C1 group. The C20 genes are all partial models (187–260 aa), so their lack of a MEROPS match should be interpreted with caution.

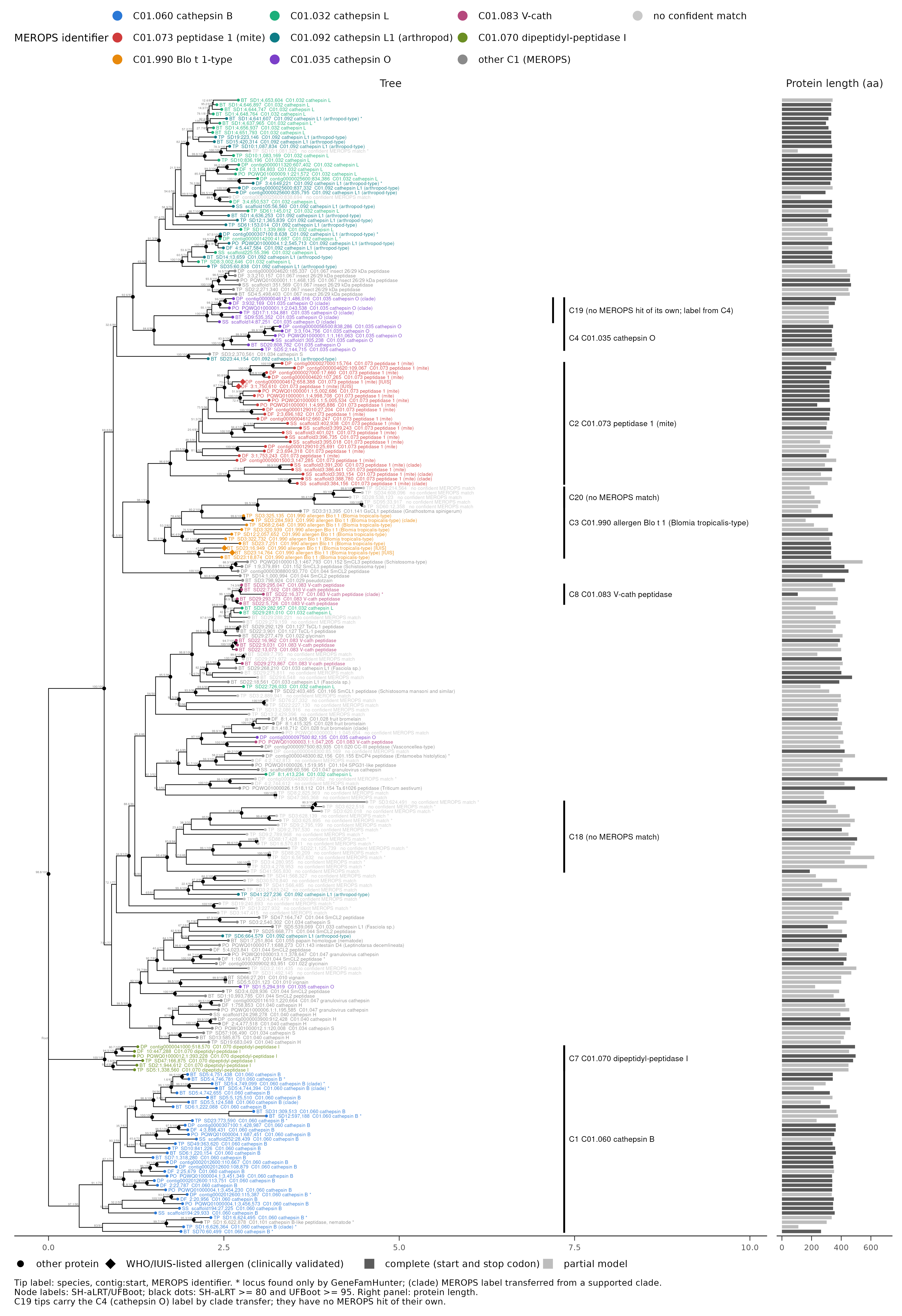

**Fig. S8.** Maximum-likelihood gene tree of the 246 GeneFamHunter C1 proteins from six mite genomes (MAFFT alignment, IQ-TREE 2, VT+R8), rooted on the cathepsin B clade. Tip labels: species (BT, B. tropicalis; DF, D. farinae; DP, D. pteronyssinus; PO, P. ovis; SS, S. scabiei; TP, T. putrescentiae), contig and start coordinate of the gene, and the final MEROPS identifier and name; (clade), label transferred from a supported clade; *, locus found only by GeneFamHunter (not in Shi et al., 2023); [IUIS] and diamond, WHO/IUIS-listed allergen (clinically validated). Tip colour: MEROPS identifier. Node labels: SH-aLRT/UFBoot; black dots, SH-aLRT ≥ 80 and UFBoot ≥ 95. Bars: MEROPS clades discussed in the text (merops_clusters.tsv); C19 (no MEROPS hit of its own; label inherited from C4) is nested within C4. Right panel: protein length; dark, complete model (start and stop codon); light, partial model. A vector version is provided as FigS8_merops_tree.pdf.

### Supplementary Note S5. Clustering and filtering parameters

Table S8. Main parameters of the GeneFamHunter clustering, ROI and filtering steps.

| **Parameter (config.env)** | **Default** | **Role** |
| --- | --- | --- |
| window / step | 1,400 / 700 bp | Overlapping genomic windows translated in six frames |
| MSR_RULE | continuity_dm | Rule set used to join or split collinear hits |
| MAX_MSR_GAP_TANDEM | 500 bp | Maximum MSR gap for joining two pieces of one domain (mite calibration: real joins ≤ 399 bp, real splits ≥ 523 bp) |
| MAX_MSR_GAP_ISOLATED | 5,000 bp | Maximum gap when the next piece continues the model (−9 to +30 positions; HMMER and rpstblastn only) |
| REDUND_MIN / JAC_MIN | 30% / 60% | Model re-read (fraction of model length / Jaccard of model ranges) that opens a new copy |
| EXT5 / EXT3 | 750 / 750 bp | Flanks added to domain MSRs to form ROIs |
| gEXT5 / gEXT3 | 360 / 180 bp | Flanks for full-length protein hits (PSI-BLAST, DELTA-BLAST) |
| CDD_EVAL | 10⁻³ | RPS-BLAST E-value for protein-level validation |
| MIN_DOMAIN_COV_IN_MODEL | 50% | Separates gene_model_missed_domain from spurious_nt_hit |
| MIN_HIT_COV | 50% | Minimum coverage of the merged-hit span by the gene model (high-quality set) |
| TREE_VERDICTS | confirmed, split fragment with target domain | Verdicts accepted into the high-quality set and gene tree |

### Supplementary Note S6. Family membership compared with orthology groups

#### Methods

*Data.* The six QfO proteomes (release 2026_02, canonical proteins) were used as in Note S2. The reference comprised the 128 canonical proteins annotated in UniProt (October 2026) with InterPro peptidase C1A (IPR000668), the family modelled by PF00112; they carry the 130 domain loci of Note S2. Bleomycin hydrolases (peptidase C1B; InterPro IPR004134, Pfam PF03051, MEROPS C01.084) are not a PF00112 target but resemble C1A enough to be found by PF00112 searches, so they were counted as a separate category; the six proteomes contain four (*D. melanogaster* X2JEJ0 and A1ZBU9, *T. castaneum* D2A4S5 and *D. magna* A0A0P5X8K8). Fused proteins have two or more PF00112 domains (HMMER 3.3.2, PF00112.30, domain i-E-value < 0.01) whose minimum search regions are > 20 residues apart, the rule GeneFamHunter uses to remove fused protein evidence (Note S1).

*Orthology groups.* OrthoFinder 3.1.5 (Emms and Kelly, 2019) was run with default settings (DIAMOND, FAMSA, FastTree, inflation 1.2) on the six canonical proteomes; every orthogroup with at least one C1A protein was kept with all its members, including single-gene orthogroups, and likewise for OMA groups. OMA groups (Altenhoff *et al.*, 2024) were taken from the OMA cross-references in the QfO ID-mapping files, restricted to canonical proteins; OMA groups contain at most one protein per species, and *D. magna* has no OMA cross-references in this release. MEROPS identifiers were assigned by BLASTP against MEROPS as in Note S4.

*Protein-level validation.* The QfO runs stopped after the ROI step, so GeneFamHunter's protein-level check (Section 2.3) was applied to the annotated UniProt proteins whose CDS overlapped each ROI, in place of GeneMark-EP+ models. For intergenic ROIs, the hit span was translated in the three frames of its strand, and stretches of ≥ 30 residues between stop codons were tested. Proteins were searched against the CDD with RPS-BLAST (E ≤ 10⁻³); an ROI was confirmed if any hit belonged to the C1 superfamily (cl23744: pfam00112, pfam03051, cd00585, cd02248, cd02619, cd02620, cd02621, cd02698, smart00645) and kept if its proteins covered ≥ 50% of the merged-hit span. Script: qfo_protcheck.py (benchmark repository).

#### Results

Every ROI that overlapped a C1A or C1B protein passed the protein-level check. These ROIs cover all 128 C1A proteins and three of the four bleomycin hydrolases; the fourth, the divergent *D. melanogaster* A1ZBU9, has no PF00112 hit and was not found by any of the four searches, which used the PF00112 profile and C1A queries. Validation removed 12 loci (Table S10): seven non-C1 proteins hit at the nucleotide level, five of them with only the cathepsin propeptide inhibitor domain (I29) and detected by PSI-BLAST or DELTA-BLAST; Q8IBZ2, a *P. falciparum* α/β-hydrolase (Pfam PF12146) with a weak PF00112 hit; and four intergenic loci without a C1 domain. Five further intergenic loci (*C. elegans* 2, *D. melanogaster* 2, *T. castaneum* 1) carry a C1 domain but no annotated gene and may be unannotated genes or pseudogenes (Fig. 1E; Table S9).

OrthoFinder placed the 128 C1A proteins in 14 orthogroups, seven with several genes and seven single-gene ones, and these orthogroups contained 35 further proteins without a C1 domain (Fig. S9): 34 *D. discoideum* carbohydrate-binding proteins in OG0000223 and one uncharacterized *D. magna* protein in OG0000203. The largest orthogroup (OG0000100, 61 proteins) combined 17 MEROPS identifiers. The bleomycin hydrolases formed their own orthogroups (OG0006906 with three and a single-gene orthogroup for A1ZBU9). OMA placed 98 C1A proteins in 84 one-to-one groups, with no other members; the other 30, including all 23 *D. magna* proteins, are in no OMA group, and the bleomycin hydrolases are in separate OMA groups.

Table S9. C1A family membership in the six QfO proteomes (Fig. 1E). UniProt C1A: reference proteins (fused in brackets). C1B: bleomycin hydrolases, counted separately. GeneFamHunter: proteins recovered after protein-level validation (+ C1 loci without an annotated gene). OrthoFinder and OMA: C1A proteins in groups that contain at least one C1A protein (+ other members, none of which has a C1 domain); no C1B protein falls in these groups.

| Species | UniProt C1A (fused) | UniProt C1B | GeneFamHunter C1A (+ loci without gene) | GeneFamHunter C1B | Removed at protein level | OrthoFinder C1A (+ other) | OMA C1A (+ other) |
| --- | --- | --- | --- | --- | --- | --- | --- |
| C. elegans | 29 (0) | 0 | 29 (+2) | 0 | 1 | 29 (+0) | 27 (+0) |
| D. melanogaster | 10 (0) | 2 | 10 (+2) | 1 | 4 | 10 (+0) | 10 (+0) |
| T. castaneum | 25 (2) | 1 | 25 (+1) | 1 | 2 | 25 (+0) | 23 (+0) |
| D. magna | 23 (0) | 1 | 23 (+0) | 1 | 2 | 23 (+1) | 0 (+0) |
| D. discoideum | 25 (0) | 0 | 25 (+0) | 0 | 1 | 25 (+34) | 25 (+0) |
| P. falciparum | 16 (0) | 0 | 16 (+0) | 0 | 2 | 16 (+0) | 13 (+0) |
| Total | 128 (2) | 4 | 128 (+5) | 3 | 12 | 128 (+35) | 98 (+0) |

Table S10. QfO loci removed or added by protein-level validation. Class: non-c1, ROI overlapping an annotated protein without a C1 hit; weak-c1, weak PF00112 hit; intergenic, no annotated gene. Best CDD hit of the protein (or of the translated hit span for intergenic loci), RPS-BLAST E ≤ 10⁻³.

| Species | Protein or ROI | Class | Best CDD hit | Outcome |
| --- | --- | --- | --- | --- |
| CAEEL | I2HAL7 | non-c1 | none | removed |
| DAPMA | A0A164HWH7 | non-c1 | Inhibitor_I29 (smart00848), E = 5e-17 | removed |
| DAPMA | LRGB01002829:116420-118735(-) | intergenic | none | removed |
| DICDI | Q54UH4 | non-c1 | CBM49 (pfam09478), E = 2e-27 | removed |
| DROME | A1Z9I0 | non-c1 (4 ROIs) | Inhibitor_I29 (pfam08246), E = 1e-13 | removed |
| DROME | A1ZBK7 | non-c1 | Inhibitor_I29 (pfam08246), E = 3e-14 | removed |
| DROME | AE013599:20484211-20486394(+) | intergenic | none | removed |
| DROME | AE014298:19288248-19290446(+) | intergenic | none | removed |
| PLAF7 | AL844501:107191-109443(-) | intergenic | PST-A (TIGR01607), E = 5e-29 | removed |
| PLAF7 | Q8IBZ2 | weak-c1 | PST-A (TIGR01607), E = 3e-140 | removed |
| TRICA | A0A139WFR3 | non-c1 | Inhibitor_I29 (smart00848), E = 3e-14 | removed |
| TRICA | D6WR24 | non-c1 | Inhibitor_I29 (smart00848), E = 2e-20 | removed |
| CAEEL | BX284604:15204900-15207101(+) | intergenic | Peptidase_C1A (cd02248), E = 4e-19 | kept (no annotated gene) |
| CAEEL | BX284605:20224115-20226313(-) | intergenic | Peptidase_C1A (cd02248), E = 1e-28 | kept (no annotated gene) |
| DROME | AE013599:12954067-12956274(-) | intergenic | Peptidase_C1A (cd02248), E = 2e-05 | kept (no annotated gene) |
| DROME | AE014297:6027213-6029425(+) | intergenic | Peptidase_C1A (cd02248), E = 5e-56 | kept (no annotated gene) |
| TRICA | CM000282:1057622-1059192(+) | intergenic | Peptidase_C1A (cd02248), E = 6e-42 | kept (no annotated gene) |

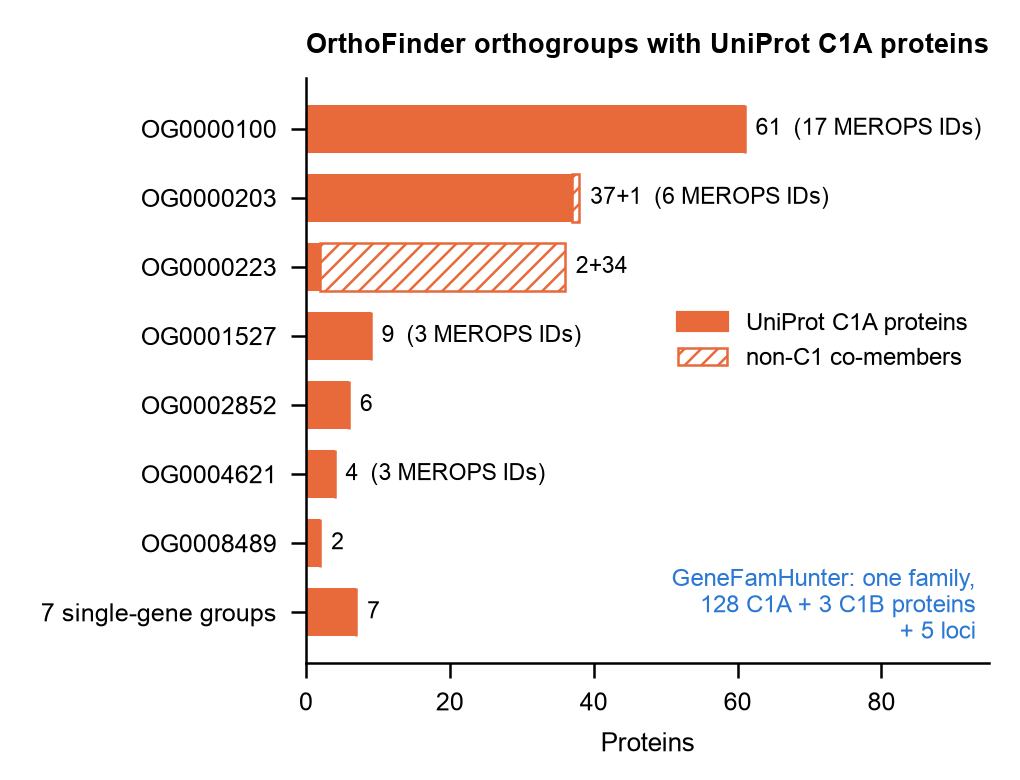

**Fig. S9.** OrthoFinder orthogroups that contain UniProt C1A proteins in the six QfO proteomes. Bars show C1A proteins (filled) and members without a C1 domain (hatched); the number of distinct MEROPS identifiers among labelled members is given where > 1. GeneFamHunter reports the family as one set: 128 C1A and three C1B proteins and five loci without an annotated gene.
